# Cx43 Enhances Response to BRAF/MEK Inhibitors by Reducing DNA Repair Capacity

**DOI:** 10.1101/2024.07.15.601645

**Authors:** Adrián Varela-Vázquez, Amanda Guitián-Caamaño, Paula Carpintero-Fernández, Vanesa Álvarez, Alexander Carneiro-Figueira, Marta Varela-Eirín, Teresa Calleja-Chuclá, Susana B Bravo-López, Anxo Vidal, Juan Sendón-Lago, Marina Rodríguez-Candela Mateos, José R Caeiro, Miguel G. Blanco, Guadalupe Sabio, María Quindós, Carmen Rivas, David Santamaría, Carlos Fernandez-Lozano, Eduardo Fonseca, Pablo Huertas, Berta Sánchez-Laorden, Constance Alabert, María D. Mayán

**Affiliations:** Instituto de Investigación Biomédica de A Coruña (INIBIC), Servizo Galego de Saúde (SERGAS), Universidade da Coruña (UDC), A Coruña, Spain; CELLCOM Research Group. Biomedical Research Center (CINBIO) and Institute of Biomedical Research of Ourense-Pontevedra-Vigo (IBI), University of Vigo. Edificio Olimpia Valencia, Campus Universitario Lagoas Marcosende, 36310, Pontevedra, Spain; Division of Molecular, Cell, and Developmental Biology, School of Life Sciences, University of Dundee, Dundee DD1 5EH, Scotland, UK; Pharmacy Service CHUAC. Instituto de Investigación Biomédica de A Coruña (INIBIC), Servizo Galego de Saúde (SERGAS), Universidade da Coruña (UDC), A Coruña, Spain; Proteomics Laboratory, Instituto de Investigación Sanitaria de Santiago de Compostela (IDIS), Complexo Hospitalario Universitario de Santiago de Compostela (CHUS), Universidade de Santiago de Compostela (USC), Santiago de Compostela, Spain; CiCLOn, Centro Singular de Investigación en Medicina Molecular y Enfermedades Crónicas (CIMUS), Universidade de Santiago de Compostela, Instituto de Investigación Sanitaria de Santiago de Compostela (IDIS), E15782, Santiago de Compostela, Spain; Department of Physiology-Center for Research in Molecular Medicine and Chronic Diseases (CIMUS), Universidade de Santiago de Compostela, Santiago de Compostela, Spain; Department of Orthopaedic Surgery and Traumatology, Complexo Hospitalario Universitario de Santiago de Compostela (CHUS), Universidade de Santiago de Compostela (USC), Santiago de Compostela, Spain; Department of Biochemistry and Molecular Biology, CIMUS, Universidade de Santiago de Compostela-Instituto de Investigación Sanitaria (IDIS), Santiago de Compostela, A Coruña 15782, Spain; Centro Nacional de Investigaciones Oncológicas C/Melchor Fernández Almagro, 3. 28029 Madrid, Spain; Medical Oncology Service. Instituto de Investigación Biomédica de A Coruña (INIBIC), Servizo Galego de Saúde (SERGAS), Universidade da Coruña (UDC), A Coruña, Spain; Centro de Investigación en Medicina Molecular y Enfermedades Crónicas (CIMUS), Instituto de Investigaciones Sanitarias (IDIS), Universidade de Santiago de Compostela, Santiago de Compostela, Spain; Departamento de Biología Celular y Molecular. Centro Nacional de Biotecnología (CNB), Consejo Superior de Investigaciones Científicas (CSIC), Madrid, Spain; Centro de Investigación del Cáncer, CSIC. University of Salamanca. Spain.; CITIC-Research Center of Information and Communication Technologies. Department of Computer Science and Information Technologies, Faculty of Computer Science. Universidade da Coruna, A Coruña 15071, Spain; Dermatology Service, Instituto de Investigación Biomédica de A Coruña (INIBIC), Servizo Galego de Saúde (SERGAS), Universidade da Coruña (UDC), A Coruña, Spain; Centro Andaluz de Biología Molecular y Medicina Regenerativa (CABIMER), Universidad de Sevilla-Consejo Superior de Investigaciones Científicas-Universidad Pablo de Olavide, Sevilla 41092, Spain; Instituto de Neurociencias, Consejo Superior de Investigaciones Científicas and Universidad Miguel Hernández, San Juan de Alicante, Spain

**Keywords:** BRAF and MEK inhibitors, targeted therapies, connexins, Cx43, DNA damage response, BRAF mutation, drug resistance, senescence

## Abstract

BRAF and MEK inhibitors (BRAF/MEKi) have radically changed the treatment landscape of advanced BRAF mutation-positive tumours. However, limited efficacy and emergence of drug resistance are major handicaps for successful treatments. Here, by using relevant preclinical models, we found that Connexin43 (Cx43), a protein that plays a role in cell-to-cell communication, increases effectiveness of BRAF/MEKi by recruiting DNA repair complexes to lamin-associated domains and promoting persistent DNA damage and cellular senescence. The nuclear compartmentalization promoted by Cx43 contributes to genome instability and synthetic lethality caused by excessive DNA damage, which could lead to a novel therapeutic approach for these tumours to overcome drug resistance. Based on these findings, we designed an innovative drug combination using small extracellular vesicles (sEVs) to deliver the full-Cx43 in combination with the BRAF/MEKi. This study reveals Cx43 as a new player on DNA repair and BRAF/MEKi response, underlining the therapeutical potential that this approach could eventually have in the clinic to overcome the limitations of current therapies and improve treatment outcomes for patients with advanced BRAF mutant tumours.

## Introduction

The mitogen-activated protein kinase (MAPK) pathway is a frequently mutated signaling cascade in cancer and comprises several signaling components that play key roles in tumorigenesis [1, 2]. A considerable number of tumours including melanoma, colon, lung, pancreas, and several forms of leukemia hold oncogenic driver mutations - such as in KRAS, HRAS, NRAS or BRAF - that promote constitutive activation of RAF-MEK-ERK MAPK pathway enhancing survival, proliferation and tumour progression [1, 3–5].The development of BRAF and MEK inhibitors (BRAF/MEKi) led to a paradigmatic change in management of BRAF-mutated melanoma, lung, and anaplastic thyroid cancers [6–8] with a major impact on the treatment of metastatic melanoma driven by the BRAF^V600E^ mutation, which represents 40-60 % of all melanoma cases [9–11]. However, the clinical benefit to this combined therapy is limited by the genetic and/or non-genetic adaptive mechanisms that lead to intrinsic or acquired resistance [12–15]. These clinical outcomes highlight the need to further explore tumour vulnerabilities to improve the efficacy and the clinical response to BRAF/MEKi.

Among the non-genetic mechanisms involved in the efficacy/resistance of these inhibitors, the senescence phenotype was suggested to drive treatment resistance to BRAF inhibitors (BRAFi) [16–18]. However, the role of senescence in BRAF/MEKi efficacy and resistant is not clearly defined. In fact and despite significant research effort and new treatment strategies, such as the combination of senolytics with BRAF/MEKi (i.e. NCT01989585 [18]), drug resistance continues to be the principal limiting factor to achieve cures in patients with BRAF^V600E^ (amino acid substitution at codon 600 (V600E)) mutated tumours. Thus, understanding the vulnerabilities of tumour cells in the context of BRAF^V600E^ mutation and therapy together with a better understanding of mechanisms involved in drug efficacy and drug resistance are critical to develop new therapeutic strategies that lead to the elimination of minimal residual disease responsible for driving therapy relapse.

Connexin43 (Cx43) is a transmembrane protein involved in cellular senescence [19] that participates in the tumour microenvironment and cell-to-cell communication via small extracellular vesicles (sEVs), tunnelling nanotubes, hemichannels and gap junctions (GJs) [20]. Cx43 does not solely exert its function via hemichannels or GJs; it also has channel-independent functions [21, 22]. Its role in cancer is context-specific [23, 24] and despite that it was reported to reduce tumour cell proliferation in certain models of melanoma (i.e. B16-BL6 cell line) [25–28], its role in BRAF-mutated tumours has not been explored yet. Given the involved pathways where Cx43 exerts a role, and due to advancements in personalized and targeted therapies, conducting cancer studies within a specific molecular context has become imperative. Hence, here we have studied the role of Cx43 in this molecular background and under BRAF/MEKi treatment. We also investigated the nuclear role of Cx43 and the transcriptional profile, connecting Cx43-induced changes to different pathways and new vulnerabilities that increase efficacy of treatments.

Our results unveil the anti-tumour activity of Cx43 in BRAF^V600E^ mutant tumours. We have identified a novel mechanism underlying the transition from a drug-sensitive to a drug-resistant phenotype, pinpointing Cx43 as a key target to overcome resistance to BRAF/MEKi. By maintaining Cx43 overexpression, tumour cells do not evade BRAF/MEKi treatment and overcome drug resistance. Further, upregulation of Cx43 in double resistant (DR) cells to BRAF/MEKi, dabrafenib and trametinib respectively, re-sensitizes DR cells to cell death. Our current research demonstrates that Cx43 engages DNA damage response and influences in DNA repair capacity at the nuclear periphery, by interacting with nuclear lamins, chromatin, and several DNA repair complexes. These Cx43-lamin-associated domains interfere with DNA repair, leading to genome instability, cellular senescence, and cell death in BRAF mutated cells, and induce synthetic lethality under BRAF/MEKi treatment. We have also developed a therapeutic mRNA/protein delivery approach using membrane-enclosed nanoparticles termed small extracellular vesicles (sEVs) for the delivery of Cx43 to target cells. Together, our research provides a strong rationale supporting the role of Cx43 in enhancing the efficacy of current targeted therapies against cancers harbouring BRAF driver mutations by exploiting a vulnerability of tumour cells in their capacity to repair DNA or survive with certain DNA damage. This approach offers a unique opportunity for cancer therapy by inducing genome instability and synthetic lethality under these treatments.

## Results

### Cx43 upregulation acts as a tumour suppressor in tumours carrying BRAF or NRAS driver mutations

While Cx43 has been proposed to reduce proliferation in specific melanoma models [20], it is important to note that melanoma exhibits high heterogeneity, characterized by various genomic subtypes [29]. Additionally, Cx43 is a transmembrane protein primarily localized within cell membranes and displays altered localization or levels in certain neoplastic cells [30]. The Cancer Cell Line Encyclopaedia database (CCLE) showed that the expression of Cx43 gene (*GJA1*, mRNA levels) is highly variable in different tumours, with relatively low expression in melanoma (red square, Fig. 1a) and other types of cancer such as colorectal cancer and several types of lymphoma (Fig. 1a). Profiling of its expression across tumour and normal samples showed that Cx43 gene expression is significantly lower in skin cutaneous melanoma (SKCM) samples compared to normal samples from the Cancer Genome Atlas (TCGA) and the Genotype-Tissue Expression Project (GTEx) (Fig. 1a). Considering these results, we decided to investigate the function of Cx43 on BRAF mutant tumour cells and BRAF mutant tumour cells transfected with a vector to upregulate Cx43 protein levels (BRAF^V600E^-Cx43) (Fig. 1b). We verified that Cx43 mRNA and protein levels were upregulated by real-time quantitative polymerase chain reaction (RT-qPCR), flow cytometry, western-blot and immunofluorescence (IF) assays (Fig. 1b and Fig. S2f). Upregulation of Cx43 restored the unmodified and phosphorylated forms of the protein (43 kDa) detected by western-blot and Cx43 clustered into plaques at sites of cell-cell contacts observed by IF (Fig. 1b). Next, Cx43 expression and protein levels were analysed in a panel of established human melanoma cell lines BRAF- and N-RAS mutants, the most common driver mutations in melanoma (Fig. S1a, b). NRAS and BRAF mutant tumour cells barely contain Cx43 protein, which was detected in the cytoplasm by IF assays with a diffuse and weak pattern (images overexposed in Fig.S1a and S1b). To confirm these findings in human melanoma samples, immunohistochemistry (IHC) was employed to detect the Cx43 staining pattern in both primary (n=19) and metastatic melanoma (n=4) tissues (see Fig. S1c). Cx43 IHC typically shows strong granular membrane and cytoplasmic localization in the different layers of the epidermis (Fig. S1d). However, the Cx43 staining in melanoma cells showed cytoplasmic localization and a diffuse and weak staining in all cases (overexposed images, Fig. S1c), including lung metastasis samples (overexposed images, Fig. S1e).

**Figure 1.**
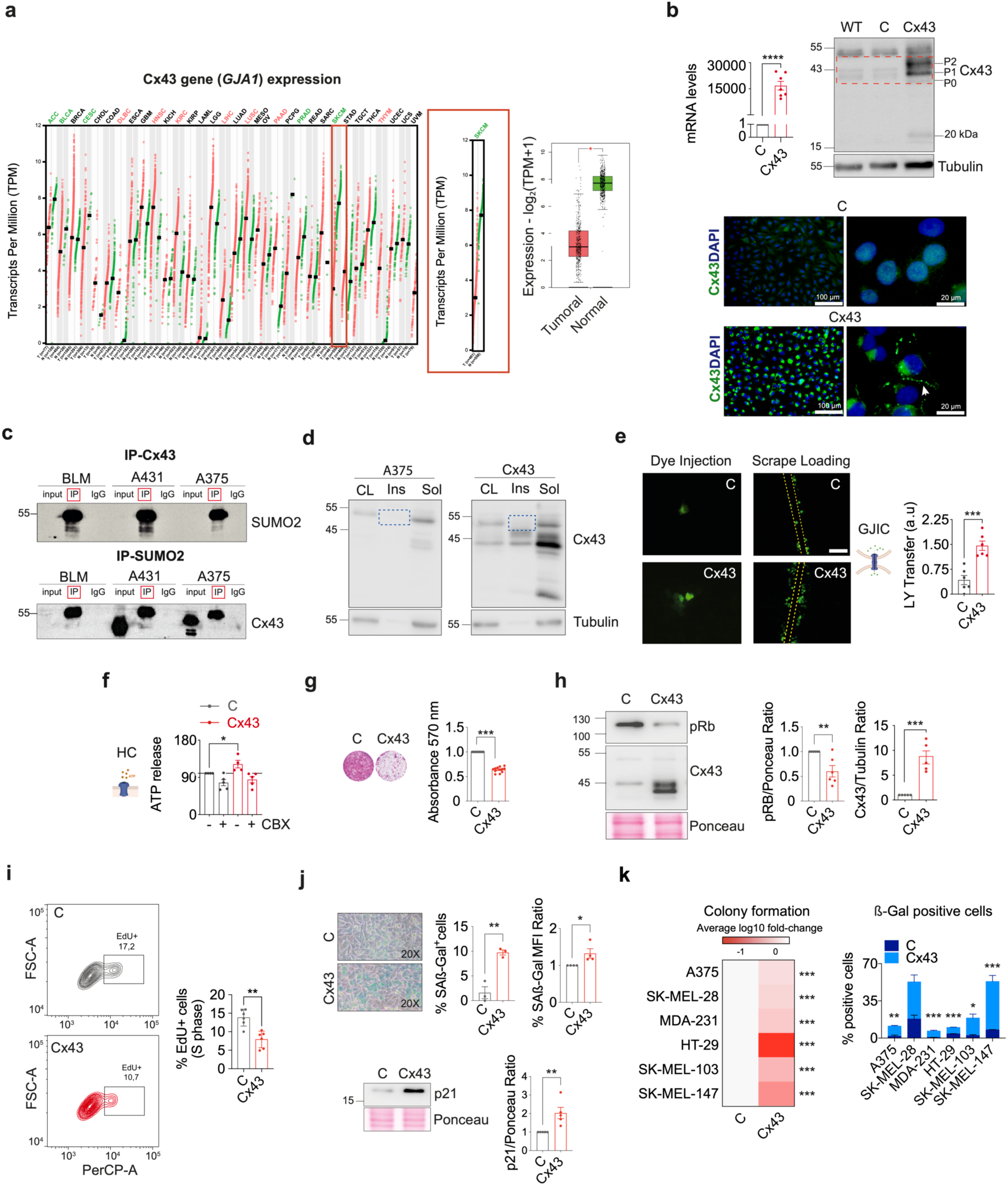
Overexpression of Cx43 decreases proliferation and increases senescence in BRAF and NRAS cancer cells. (a) Box plots of GJA1 gene expression across different tumour types (CCLE dataset). Image from http://www.broadinstitute.org/ccle/home. (GJA1 gene expression profile across all tumour samples and paired normal tissues. Log Scale TPM or log2 (TPM+1) transformed expression data is showed for plotting. Images from GEPIA (cancer-pku.cn). (b) Cx43 RT-qPCR (n= 3), Cx43 immunoblot and Cx43 staining of A375 cells and after Cx43 overexpression (WT, non-transformed cell line; C, transformed with empty vector; and Cx43, transformed with Cx43 vector). White arrows show the localization of Cx43 in the membrane. Scale bars: 100 µm (Cx43 and DAPI). Arrows indicates the non-phosphorylated (P_0_) and phosphorylated (P_1_ and P_2_) Cx43 isoforms. Tubulin was used as a loading control. (c) Immunoprecipitation (IP) and immunoblotting of Cx43 and SUMO2 in A375, BLM and A451 cell lines. (d) Immunoblotting for Cx43 cellular localization in total cell lysate (CL), insoluble (Ins) and soluble (Sol) fractions in A375 and A375 with Cx43 overexpressed (A375-Cx43) cells. Tubulin was used as loading control. (e) Images and quantification of dye transfer (left) and scrape loading (right, n= 6) assays in tumour cells A375-C or A375-Cx43 cell lines. (f) Quantification of ATP release in A375-C and A375-Cx43 cells in basal conditions and treated with 100 µM of the hemichannel blocker, carbenoxolone (CBX). (g) Images and quantification of colony formation assays in A375-C and A375-Cx43 cells for 7 days (n= 3, 5 replicates each). (h) Immunoblotting and quantification for pRB and Cx43 in A375-C and A375-Cx43 cells. Ponceau staining was used as loading control. (i) Cell cycle flow cytometry analysis and quantification of phase S in A375-C and A375-Cx43 cells. (n= 3, 2 replicates each). (j) Representative images of SAß-Galactosidase activity and quantification in A375-C and A375-Cx43 cells (n= 4). Quantification of SAß-Galactosidase detected by flow cytometry (MFI) (n= 5). Immunoblotting and quantification of senescence marker p21 (n= 3). Ponceau was used as loading control. (k) Heatmap showing the colony formation capacity of different BRAF and NRAS mutant cell lines. The data were normalized to C cells. SAß-Galactosidase quantification (% of positive cells) in different BRAF and NRAS cell lines with the Cx43 overexpression (n=3). Mann-Whitney or two-tailed Student’s t-test were used to calculate the significance represented as followed: ns: no significant, * p < 0.05, ** p < 0.01. *** p < 0.001. Data is presented as mean ± SEM

On the other hand, when Cx43 protein levels were analysed by western-blot, we were surprised to find that BRAF and NRAS mutant cells contain a different Cx43 band pattern (images overexposed, Fig. S1b). Both, BRAF and NRAS mutant cells mainly contain high molecular weight bands (up to 43 kDa), exhibiting lower levels of the bands corresponding to the unmodified and phosphorylated forms of the protein (43 kDa) (Fig. S2b and Fig. S1b and Fib. 1b for validating antibody specificity and phosphorylated forms). We decided to investigate whether the main band detected under BRAF gene mutation (around 50-55 kDa) corresponded to post-transcriptional modifications involved in protein stability or degradation. However, no changes were observed by western-blot when BRAF mutant cells were treated with 10 µM of proteasome inhibitor MG132 for 24 hours (h) to detect Cx43 degradation (Fig. S2c). As Cx43 has previously been reported to be modified by the small ubiquitin-related modifier (SUMO) [31, 32], immunoprecipitation (IP) experiments were performed in different cell lines to determine whether the higher molecular weight band corresponds to the SUMOylated form of the protein (Fig. 1c). The immunoprecipitated proteins analysed by western-blot and using anti-Cx43 and anti-SUMO2 antibodies showed that the 55 kDa band corresponded to the SUMOylated form of Cx43 (Fig. 1c). Indeed, when tumour cells were treated with different concentrations of the SUMOylation inhibitor ginkgolic acid (GA) for 48 h, we observed a decrease in the “global” Cx43 levels and also in the 55 kDa band (Fig. S2d), suggesting that SUMOylation may play a role in Cx43 stability. Finally, the treatment of BRAF mutant cells with 3.5 µM of the proteasome inhibitor bortezomib (BTZ) revealed a modest increase in Cx43 protein levels after 6-8 h of treatment, but we did not detect significant changes in the protein bands pattern (Fig. S2e).

Analysis of the insoluble cell membrane fraction by western-blot confirmed the presence of Cx43 mainly in the soluble cytoplasmic cell fraction in controls (A375 empty vector-transfected cells (C)) and the presence of Cx43 in cytoplasmic (soluble) and membrane (insoluble) fractions in A375 cells transfected with the Cx43 vector (Fig. 1d). As showed before, these results indicated that ectopic upregulation of Cx43 in BRAF mutant tumour cells, restored Cx43 membrane localization (Fig. 1b, IF). However, we noted that the band corresponding to the SUMOylated form of Cx43 was not found in the insoluble membrane fraction (blue dashed squares, Fig. 1d), indicating that Cx43 is not found SUMOylated at the membrane, at least in tumour cells carrying an activating mutation in the BRAF kinase. Then, we aimed to measure how Cx43 restoration at the membrane of BRAF-mutant cells affects the activity of GJs and hemichannels. First, we performed dye injection and scrape loading assays. Cx43 upregulation restored direct cell-to-cell communication, mediated by GJs between coupled BRAF mutant cells, as detected by lucifer yellow/dye transfer analysis (Fig. 1e). In addition, upregulation of Cx43 enhanced the hemichannel activity in these tumour cells, as measured by ATP release and in the presence of the channel (GJs and hemichannel) blocker carbenoxolone (CBX) (100 µM for 30 minutes (min)) (Fig. 1f).

Regarding the association of *GJA1* with the mutational status of the oncoproteins involved in the RAS/MAPK pathway, slightly higher levels of *GJA1* were found in BRAF melanoma mutants (pooled cases and primary melanoma lesions), whereas slightly lower levels of GJA1 were characteristic of NRAS melanoma mutants (pooled cases and melanoma metastasis), according to the TIMER database (v. 2; http://timer.cistrome.org/). No significant association was found between GJA1 levels and KRAS melanoma mutants (Fig. S1f). The CCLE and NCI-60 database showed a modification rate for Cx43 gene (mutation, amplification, deep deletion, or multiple alterations) of 6.85 % in the 73 cases of melanoma analysed (Fig. S1g). According to TCGA, *GJA1* modifications represent only 5.19 % of the 443 cases analysed. However, these modifications did not affect Cx43 gene expression analysed by the Illumina RNASeq (log_2_) system in the altered and unaltered groups (Fig. S1i).

Interestingly, upregulation of Cx43 restored its channel activity and decreased the ability of BRAF^V600E^ cell lines to form colonies (Fig. 1g). Treatment with 100 µM of CBX suggested that the reduction in cell proliferation under Cx43 upregulation occurred in a channel-independent manner (Fig. S2g). The effect of Cx43 upregulation on cell cycle progression was confirmed by a decrease in phospho-Rb+ (pRB) by western-blot (Fig. 1h) and by assessing cell cycle progression by quantifying the amount of 5-ethynyl-2’-deoxyuridine (EdU) incorporated into DNA by Fluorescence-activated cell sorting (FACs) (Fig. 1i). Data reported by our group demonstrate that Cx43 promotes a stable state of growth arrest known as cellular senescence [33, 34]. Indeed, the cell cycle delay observed in BRAF^V600E^-Cx43, correlated with an increase in the senescence-associated ß-galactosidase (SA-ß-gal) activity quantified by microscopy and flow cytometry (Fig. 1j) and with a significant increase in the levels of the senescence markers p21Waf1/Cip1 (Fig. 1j), p16INK4a (p16) and p53 (Fig. S3a). Senescence program involves a secretory activity known as the Senescence Associated Secretory Phenotype (SASP) with impact on the tumour microenvironment [35, 36]. Interestingly, BRAF^V600E^-Cx43 showed a reduction in the levels of a some secreted SASP factors in comparison with control cells, including chemokines associated with poor survival in cancer such as CCL20, IP-10 [37] or angiogenin [38] (Fig. S3b). Secreted PDGF-AA, which was also downregulated in BRAF^V600E^-Cx43 cells, promotes the production of extracellular cell matrix (ECM) components by tumour-associated fibroblasts, which in turn stimulate the tumour proliferation in a paracrine manner [39]. Consistently, Cx43 upregulation in a panel of different NRAS and BRAF^V600E^ mutant human tumour cell lines from melanoma, colon and breast cancer reduced the capacity to form colonies and enhanced cellular senescence (Fig 1k and S3c-g).

We then confirmed the anti-tumour effect of Cx43 upregulation on tumour growth *in vivo*. BRAF^V600E^ mutant cells or cells overexpressing Cx43 (BRAF^V600E^-Cx43) were subcutaneously implanted into nude mice (Fig. 2a). Upregulation of Cx43 significantly reduced primary tumour growth by 82,15 % on average (Fig. 2a). No effect on body weight was observed (Fig. S4a). IHC studies confirmed an increase in Cx43 protein levels in A375-Cx43 cell-derived xenografts and a significant decrease in cell proliferation as detected by Ki67 staining and a reduction in tumour cellularity (Fig. 2b). As Cx43 has previously been reported to induce cell death by apoptosis [26, 40–43], we also studied the mRNA levels of caspase-3 (CASP3), which was increased in BRAF^V600E^-Cx43 tumour cells in tumour sections (Fig. 2c). In fact, Cx43 increased both, cell death by apoptosis mediated by caspase-3 in 2D monocultures and cell death by apoptosis and necrosis detected by flow cytometry (Fig. 2d and Fig. S4b). Taken together, the results showed a dual antitumour function of Cx43 in BRAF^V600E^ tumour cells by promoting senescence and cell death.

**Figure 2.**
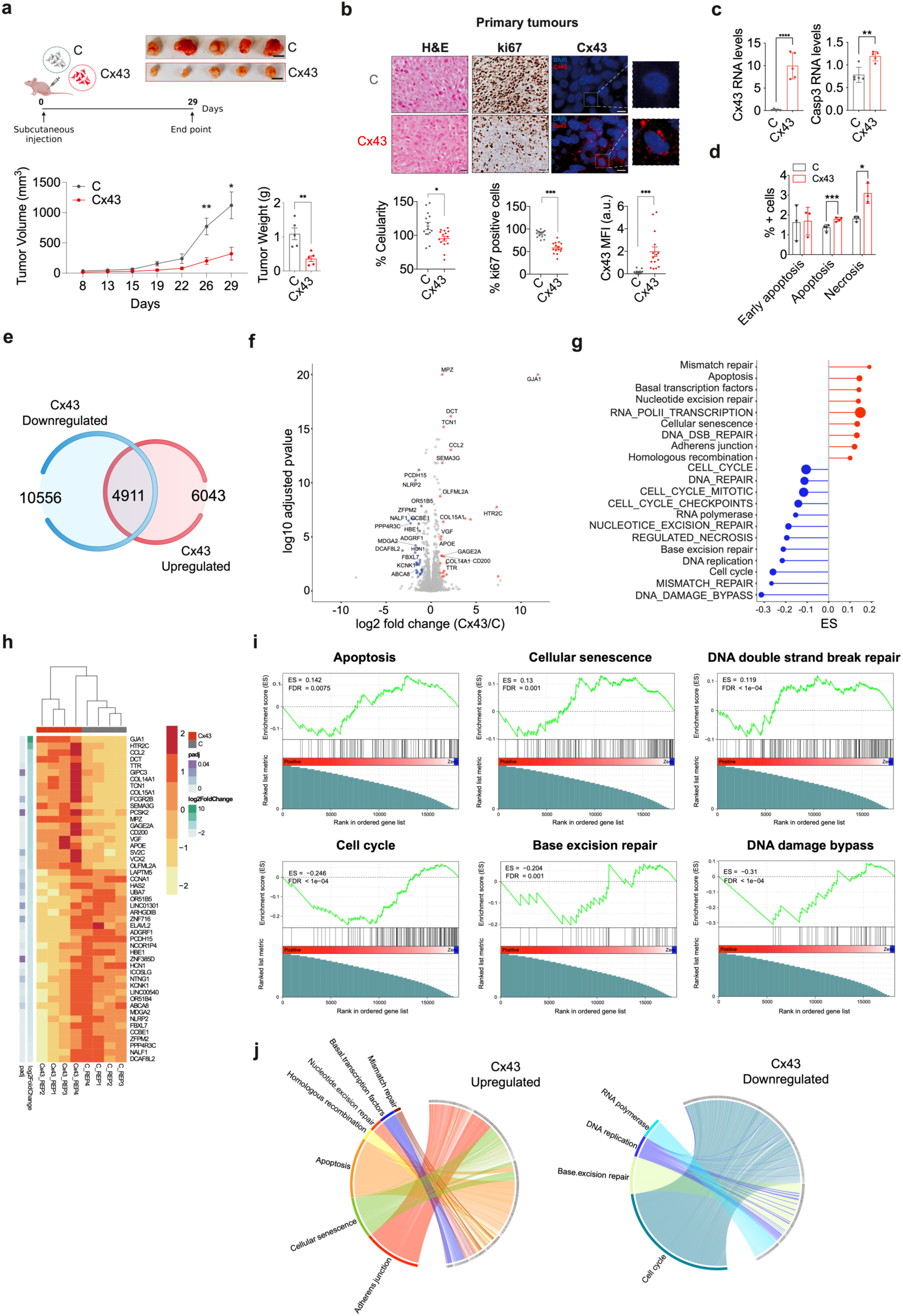
Cx43 enhances senescence, cell death and DNA damage response in BRAF cancer cells. (a) Schematic representation of the experiment workflow and images of the tumours (upper panel). Tumour growth of A375-C and A375-Cx43 xenografts in nude mice (bottom panel). Tumour weight at endpoint (n = 5 mice/group) is shown on the right. (b) Immunostaining and quantification of Cx43 and Ki67 in all tumours. Mice tumour slides were stained with haematoxylin and eosin (H&E) to measure the percentage of cellularity. Data were expressed as MFI arbitrary units (a.u.). (c) Cx43 and CASP3 mRNA levels from end point mice-derived tumours. (d) Percentage (%) of necrotic and apoptotic cells in A375-C and A375-Cx43 population cells by flow cytometry (double staining PI/YO-PRO) (n= 3-4). (e) RNA-seq analysis was performed on four replicates of A375-C and A375-Cx43. The Venn diagram shows the differentially and commonly expressed proteins in A375-C and A375-Cx43 cells. The volcano plot in (f) shows the differentially regulated transcriptomics when comparing control and Cx43 overexpressing A375 cells (LFC ≥ 1 and ≤ -1; log_10_(p-value) < 0.05). Signature proteins overexpressed in A375-C are marked in blue, and signature proteins overexpressed in A375-Cx43 are marked in red. Please refer to Table 1 for more information. (g) Pathway analysis of differentially expressed proteins was conducted using KEGG and Reactome (uppercase). The size of the circles represents the number of genes in the Leading Edge Number. Positive enrichment scores (ES) (red) indicate overexpressed pathways in A375-Cx43, while negative ES (blue) indicate downregulated pathways in A375 Cx43. Statistical significance was determined by Benjamini-Hochberg FDR adjusted p-value < 0.05. (h) A heatmap was used to compare differentially expressed genes (DEGs) between A375-C and A375-Cx43. The DEGs were ranked based on their LFC. Only DEGs with an adjusted p-value < 0.05 and |LFC| >=1 were included. Gene set enrichment analysis (GSEA) revealed positive enrichment of apoptosis, cellular senescence, and double-strand break repair signatures, and negative enrichment of cell cycle, base excision repair, and DNA damage bypass signatures in A375-Cx43 cells. (i) Chord diagrams were used to represent pathway-associated gene sets that were linked to categories of up- and down-regulated pathways in A375-Cx43 cells. The significance of each pathway was determined based on the LFC of the gene. The enrichment of genes with adjusted p-value < 0.05 and |LFC| >= 1 was studied. The cumulative value of the pathway was calculated as the sum of the LFC of the genes that form the pathway. Two-tailed Student’s t-test were used to calculate the significance represented as followed: ns: no significant, * p < 0.05, ** p < 0.01. *** p < 0.001. Data is presented as mean ± SEM.

**Table 1.**
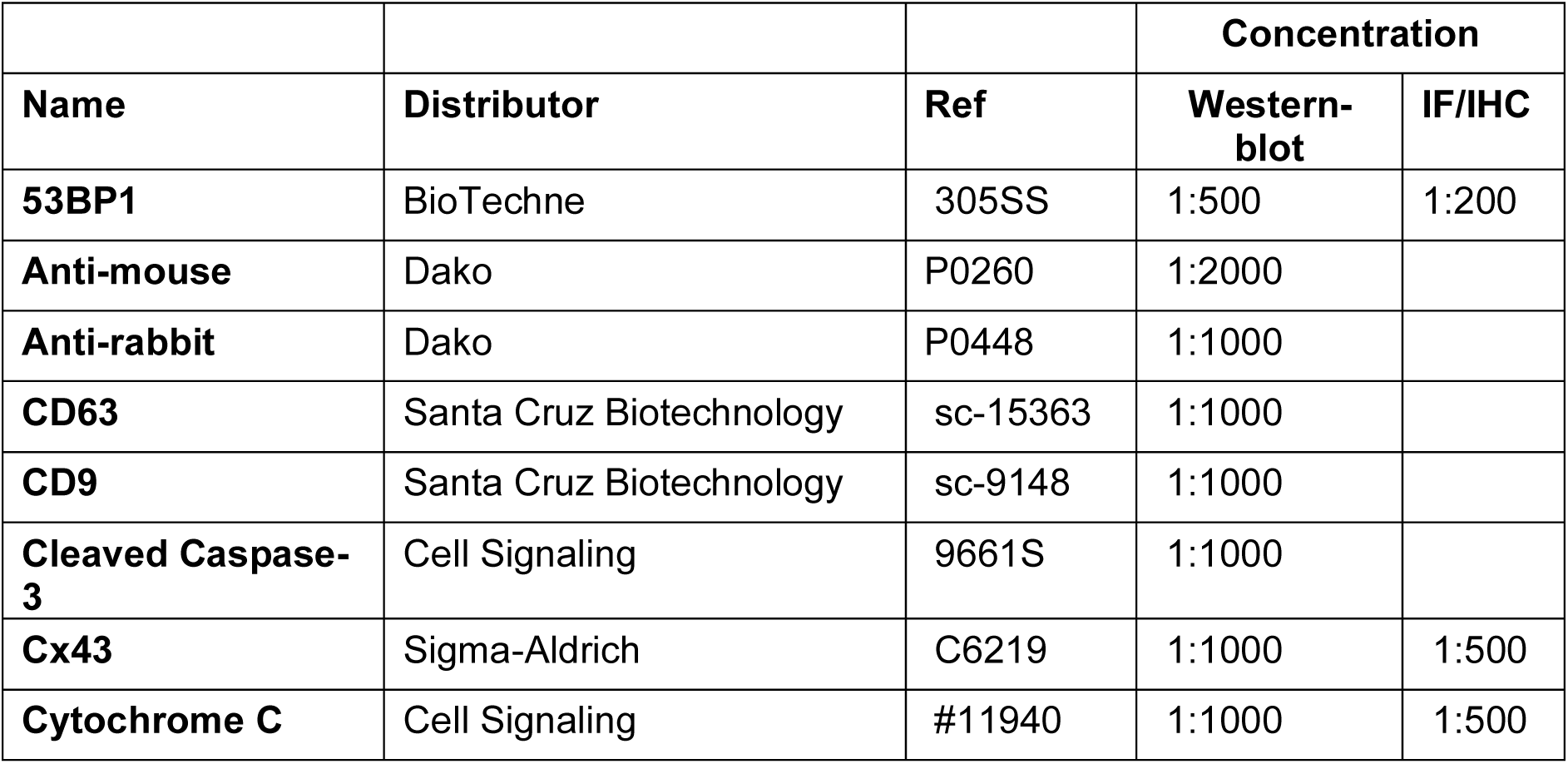
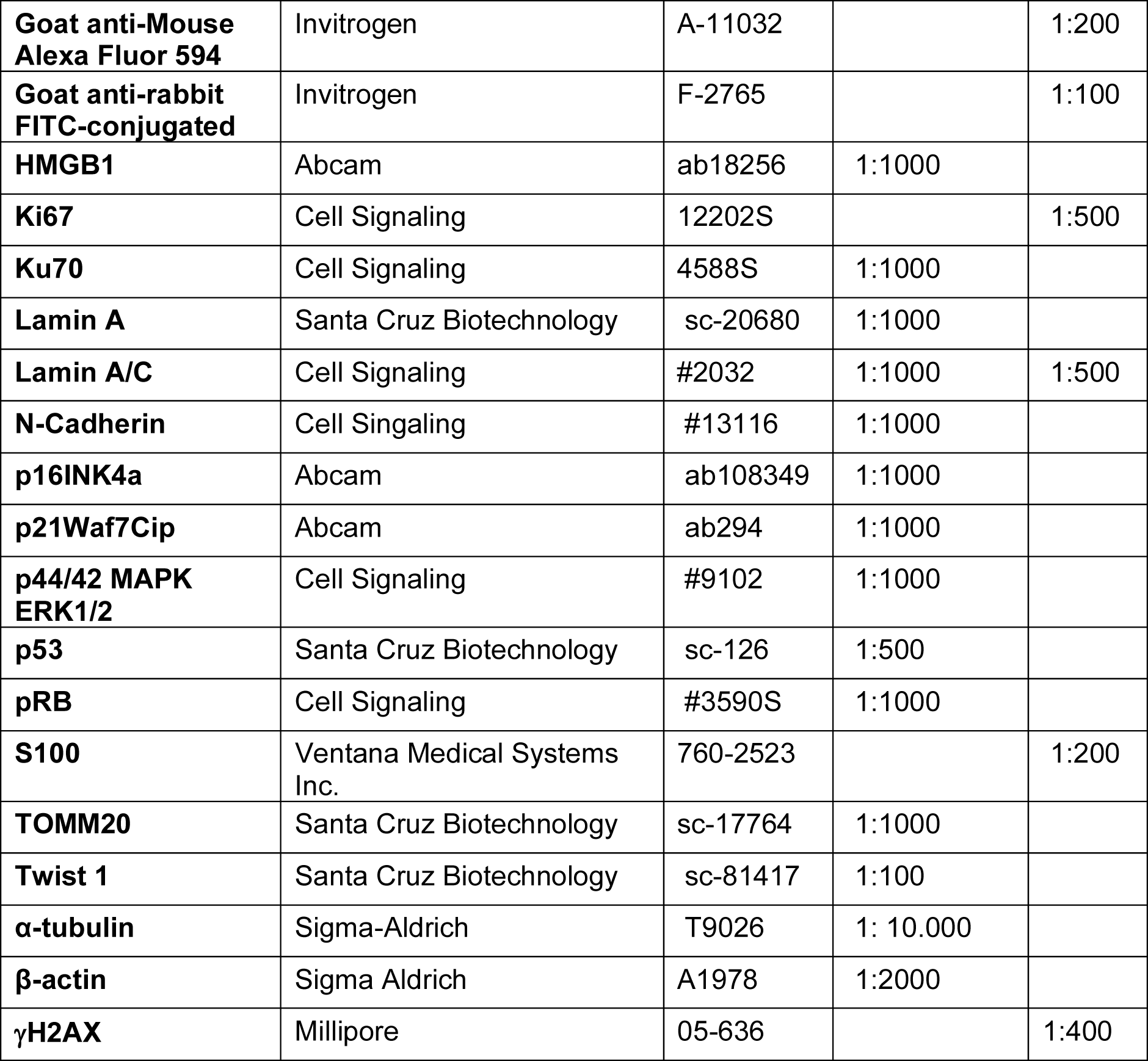
Antibodies used in this study.

**Table 2.** Cell lines.

| **Name** | **Source** | **Catalog number** |
| --- | --- | --- |
| **PCS-200** | ATCC® | PCS-200-013TM |
| **A375** | ATCC® | CRL-1619TM |
| **MDA-MB-231** | ATCC® | HTB-26TM |
| **HT-29** | ATCC® | HTB-38TM |
| **HEK-293** | ATCC® | CRL-1573TM |
| **Sk-Mel-28** | ATCC® | HTB-72TM |
| **Sk-Mel-103** |  |  |
| **Sk-Mel-147** | Sigma | SCC440 |
| **BLM** | Donated by Oncology group (INIBIC) |  |
| **MCF7** | ATCC® |  |

**Table 3.**
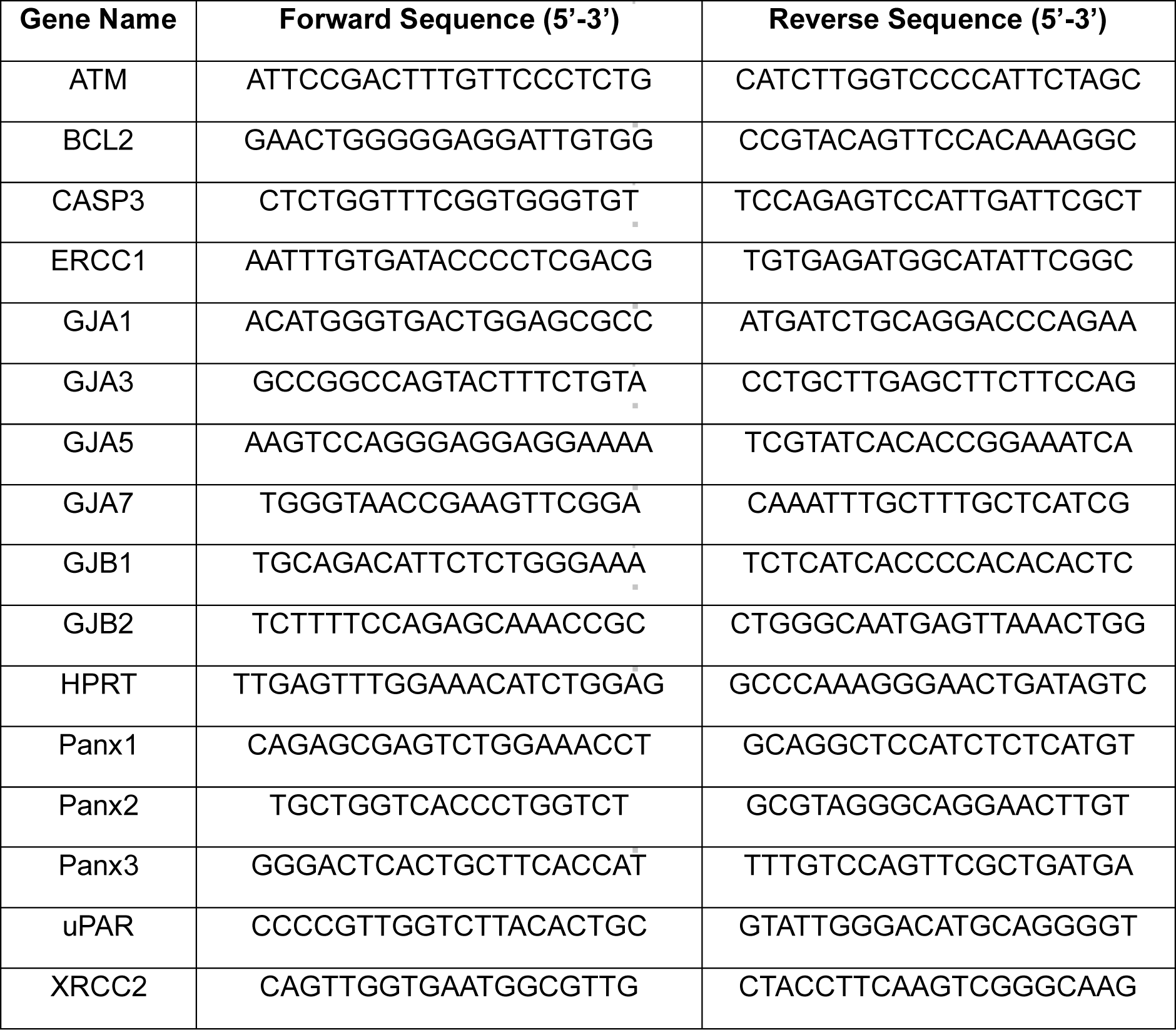
Primers.

To elucidate the mechanism by which Cx43 reduces cell cycle progression and promotes cell death, we further analysed the genes and molecular pathways modulated by Cx43 using RNA-Sequencing (Seq) in BRAF^V600E^ tumour cells. This approach allowed us to study the gene signatures and gain insight into the underlying molecular mechanisms (Fig. 2e-j). We identified 16,599 differentially expressed genes (DEGs) between control and overexpressing Cx43 cells (Fig. 2e-f). Enrichment analysis revealed the differential regulation of several pathways, particularly those associated with DNA repair and the DNA damage response (DDR) (Fig. 2g). Senescence and apoptosis-related genes were upregulated, while cell cycle pathways were downregulated in BRAF^V600E^-Cx43 cells (Fig. 2g). Surprisingly, we observed changes and an important representation of genes involved in DDR, such as an upregulation of nucleotide excision repair, DNA double-strand break (DSB) repair and homologous recombination processes, and a downregulation of base excision repair and DNA damage bypass (Fig. 2g). We have identified a set of downregulated genes in BRAF^V600E^-Cx43 cells, which are mainly involved in cell cycle progression (e.g. CCNA1, LAPTM5, HCN1), DNA damage bypass (e.g. UBA7, DCAF8L2), apoptosis inhibition (e.g. HAS2, ELAVL2), treatment resistance (e.g. HBE1) and cancer progression (e.g. NTNG1, KCNK1, ZFPM2), among others (Fig. 2h). GSEA analysis further confirmed that BRAF^V600E^-Cx43 cells showed a significant enrichment in apoptosis, cellular senescence and DNA double-strand break repair; and a down-regulation in cell cycle, base excision repair and DNA damage bypass pathways (Fig. 2i-j).

### Cx43 interacts with the nuclear lamin and interferes with the DNA damage response in BRAF^V600E^ mutant tumours

The transcriptomic data highlighted the involvement of Cx43 in DNA damage and repair processes. We therefore decided to investigate the levels of reactive oxygen species (ROS) and the DDR signalling pathway to determine the mechanism underlying the ability of Cx43 to alter cell cycle progression and to enhance cell death (Fig.3). While BRAF^V600E^-Cx43 cells contained slightly higher levels of intracellular ROS (Fig. 3a), these tumour cells have significantly increased DNA damage, as evidenced by an increase in the number the nuclear foci containing key DDR proteins, such as the adaptor protein tumour suppressor p53-binding protein 1 (53BP1), the phosphorylated serine-protein kinase ATM (pATM) and phosphorylation of H2AX at the Ser139 (γ-H2AX) (Fig. 3b). We also found that BRAF^V600E^-Cx43 cells showed upregulation of the expression of genes involved in DNA repair processes such as the DDR checkpoint kinase ATM and ERCC1, XRCC2 and uPAR, compared to control cells (Fig. 3c), supporting the hypothesis that Cx43 presence may induce persistent DNA damage foci or limit the DDR capacity. The enhanced accumulation of DNA breaks in BRAF^V600E^-Cx43 cells was confirmed by alkaline comet assay (single-cell gel electrophoresis) (Fig. 3d). Cx43 upregulation increases the comet tail-like structure corresponding to DNA damage (a mixture of single-stranded DNA [ssDNA] arising from double-strand breaks [DSBs] and single-strand breaks [SSBs]) migrating out of the nuclei in a BRAF^V600E^ mutation background (single cell, Fig. 3d). In addition, when the amount of DNA damage was increased by exposing tumour cells to 5 µM of bleomycin for 72 h, we observed a greater increase in the percentage of comet-like cells in BRAF^V600E^-Cx43 cells than in control cells (Fig. 3d). Furthermore, the results indicated that after 72 h of recovery (without bleomycin), BRAF^V600E^-Cx43 cells exhibited a significant amount of persistent DNA damage, which was partially reversed in control cells (Fig. 3d). This suggested that Cx43-expressing cells generate more DNA damage and are less efficient at repairing it compared to the control cells.

**Figure 3.**
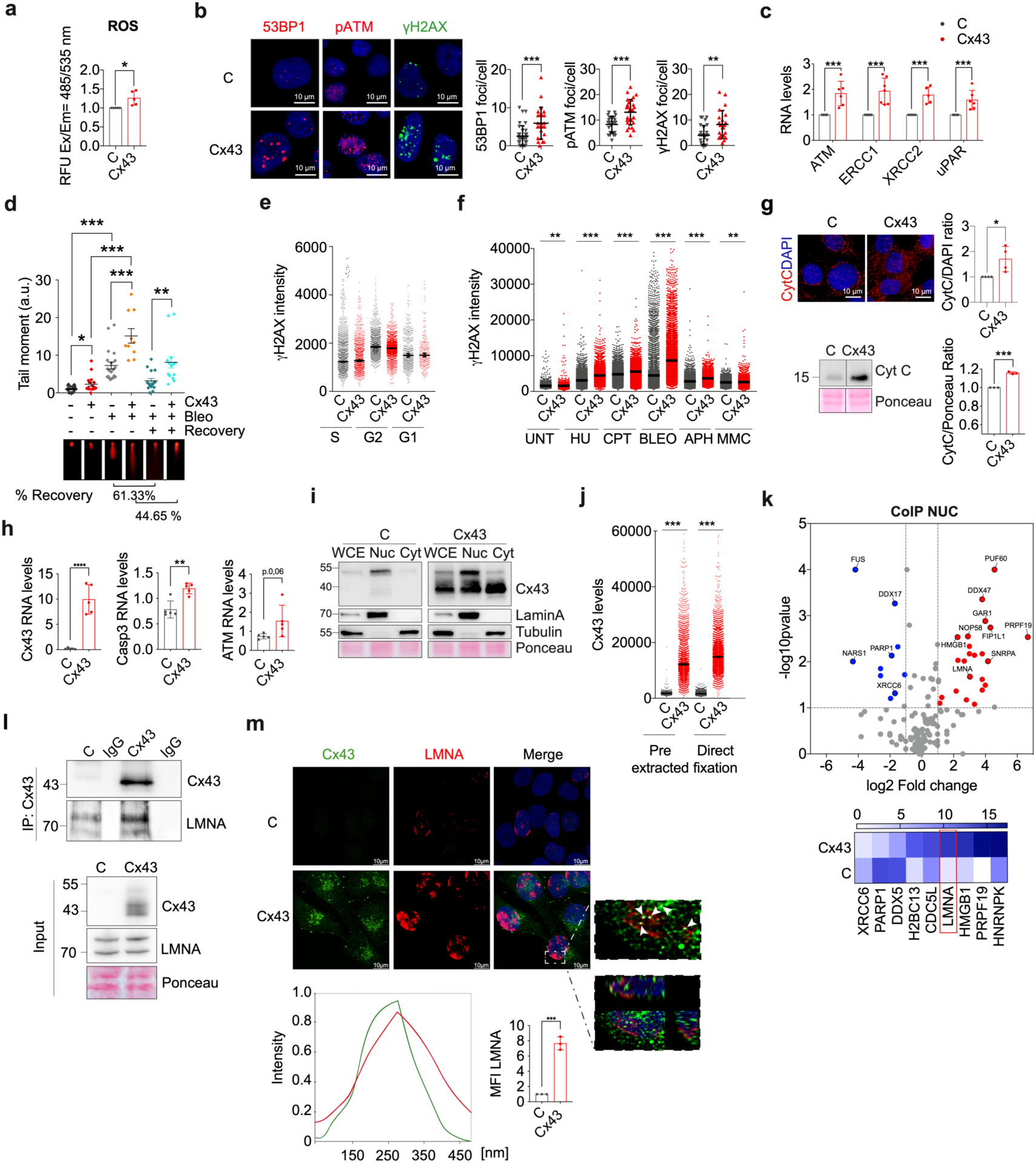
Cx43 interacts with the nuclear lamin and interferes with the DNA damage response in BRAF^V600E^ mutant tumours. (a) Cytoplasmic ROS levels in A375 cells showed a significant increase in A375-Cx43 compared to C cells. Data is represented as Relative Fluorescence Units (RFU) measured at Ex/Em= 485/535 nm. (n=5). **(b)** Representative nuclei images and quantification of the DNA damage markers 53BP1, pATM and γ -H2AX foci in A375-C and A375-Cx43 cells. **(c)** RNA expression levels of the DNA repair markers (ATM, ERCC1, XRCC2, uPAR) in A375-C and A375-Cx43 cells (n= 4-7). **(d)** Representative images from the comet assay analysis to evaluate DNA damage in C and Cx43 overexpressing cells. As a positive control for DNA damage, cells were treated with bleomycin (BLEO) 5 µM for 72 h. In addition, a control where bleomycin was removed 72 h before the analysis was used to evaluate the DNA recovery potential of the cells. Tail moment was measured with the OpenComet plug-in from ImageJ software. **(e)** Quantification of the γ-H2AX mean fluorescence intensity per cell in Cx43-overexpressing MCF7 cells (Cx43, red dots) and MCF7-C cells transfected with an empty vector (C, black dots). Cells are classified according to the cell cycle phase they belong, determined by EdU incorporation and DAPI total intensity. A pre-extraction step was included in the IF protocol. **(f)** γ-H2AX mean fluorescence intensity per cell in Cx43-overexpressing MCF7 cells (Cx43, red dots) and MCF7 control cells transfected with an empty vector (C, black dots) upon 2 h treatments with 3 mM hydroxyurea (HU), 1 µM camptothecin (CPT), 25 µg/mL BLEO, 25 µg/mL aphidicolin (APH) and 25 µM mitomycin-C (MMC). A pre-extraction step was included in the IF protocol. **(g)** Immunofluorescence of cytochrome c (CytC) release in A375-C and A375-Cx43 cells; quantification of n=3 is shown on the right. Below, protein levels of CytC were measured by western-blot in A375-C and A375-Cx43 cells; Ponceau staining was used as a loading control and quantification of n=3 is shown on the right. **(h)** Cx43, CASP3 and ATM mRNA levels from untreated mice-derived tumours. **(i)** Cx43 protein analysis in enriched nuclear (Nuc) and cytoplasmic (Cyt) fractions in A375-C and A375-Cx43 cells. Lamin A was used as control of nuclear fraction and tubulin was used as cytoplasmic fraction control. **(j)** Cx43 mean intensity per cell detected in the nuclear area in MCF7-C cells (black dots) and Cx43-overexpressing MCF7 cells (Cx43, red dots). Detection was performed in chromatin fraction (pre-extracted) or in nuclear fraction (direct fixation). **(k)** Cx43 IP was performed in the nuclear fraction (NUC) of either A375-C or Cx43 cells, and protein content was analyzed by mass spectrometry (TripleTOF 6600 System). Volcano plot (VolcaNoseR) illustrating the spectral count fold change and its significance in the common proteins shared by A375-C and A375-Cx43. Overrepresented proteins in A375-Cx43 are displayed as red dots, while blue dots represent proteins that are increased in the A375-C group. On the right, the heatmap graph represents the fold change of spectral count values and the −10log p-value of proteins involved in DNA repair processes. (**l**) Cx43 IP NUC of A375-C and Cx43 cells. Input is shown on the left, while IP is shown at the right. IgG is indicated as a negative control. **(m)** Immunofluorescence for Lamin-A/C (LMNA, in green) and Cx43 (in red) in A375-C and A375-Cx43 cells. Imaging was visualized in a confocal microscope and colocalization between LMNA and Cx43 was calculated based on Pearson’s correlation coefficient. Below, LMNA protein levels were measured by western-blot. All experiments were performed at least three independent times. Data are presented as mean ± SEM. Two-tailed Student’s t-test and one-way ANOVA were used to calculate the significance represented as followed: * P < 0.05, **P < 0.01, ***P < 0.0001.

In light with the above observations, we investigated whether the accumulation of DNA damage mediated by Cx43 impacts on genome stability in a different background and using single cell IF analysis (Fig. 3e). Overexpression of Cx43 in the MCF7 cell line did not significantly increase DNA damage at any stage of the cell cycle as measured by γ-H2AX intensity (Fig. 3e) and seems to slightly decrease when we analyze the bulk of cells in different cell cycle stages (Fig. 3f). However, when MCF7 cells were treated for 2 h with various insults that induce DNA damage by different mechanisms, including hydroxyurea (HU), camptothecin (CPT), bleomycin (BLEO), aphidicolin (APH) or mitomycin C (MMC), we observed an increase in γ-H2AX intensity in MCF7-Cx43 cells (Fig. 3f). These results suggest that following an insult that increases DNA damage, such as BRAF activation or bleomycin, Cx43 increases susceptibility to DNA damage, which can lead to senescence and to secondary tumour cell death, possibly by disrupting the cell proficiency in repairing damaged DNA. It has been described that cytochrome c (CytC) release activates cell death by apoptosis under conditions of DNA damage [44]. IF and western-blot analyses examining CytC release from mitochondria following upregulation of Cx43 in BRAF^V600E^ mutant cells confirmed the increased levels of CytC release in BRAF^V600E^-Cx43 cells (Fig. 3g). These results were confirmed in the analysis of RNA of the xenografts-derived tumours from BRAF^V600E^-Cx43 cells, that showed increased mRNA levels of ATM and CASP3 (Fig. 3h).

To identify potential factors involved in the Cx43-mediated impairment of DDR, we analysed the nuclear interactome of Cx43 by mass spectrometry. First, we studied the levels and banding patterns of Cx43 in the nucleus by western blot (Fig. 3i). Control cells contain the band corresponding to the SUMOylated Cx43 mainly in nuclear extracts and at lower levels in the cytoplasm. BRAF^V600E^-Cx43 cells contain the SUMOylated form mostly in the nuclear extract and 43 kDa bands in both cellular extracts (Fig. 3i). Since we have previously predicted different DNA-binding sequences in the Cx43 sequence [45] and we have also identified two potential nuclear localization signals (NLSs) on the C-terminal domain of Cx43 [46], we decided to investigate the binding of Cx43 to chromatin in nuclear extracts after treatment with triton to remove proteins not loaded on chromatin and after direct fixation (Fig 3j). Analysis of the pre-extracted data from individual cells confirmed the strong direct binding of Cx43 to the chromatin. Images were taken in ScanR, a high content microscope (Fig. 3j).

Label-free proteomics after pull-down of Cx43 from the nuclear fraction (Fig. 3k), revealed several known Cx43-binding proteins including several histones, HMGB1 and nuclear lamins [46] (see Table S1 and S5a). Gene Ontology (GO) analysis of cellular components identified 215 interactors within intracellular membrane-bound organelles and 87 proteins within the nucleoplasm in BRAF^V600E^-Cx43 nuclear extracts. We have identified several interactors involved in DDR, chromatin architectural proteins, transcriptional regulation and more significant multiple proteins involved in DNA repair and RNA binding and processes (Table S1). Our analysis identified the lamin-A/C (LMNA), the high mobility group protein B1 (HMGB1) and the pre-mRNA-processing factor 19 (PRP19) among the strong Cx43-associated proteins (Fig. 3k). Interestingly, these three factors are involved in DDR and chromatin remodelling processes. Other strong interactors include the nucleolar protein 56 (NOP56) required for the box C/D snoRNAs, the histone cluster 1 H2B family member L (H2BC14 involved in senescence and DNA repair and the RNA binding protein 25 (RBM25) or SART1 involved in mRNA splicing. Interestingly PRP19 has a direct role in both DDR and the prevention of transcription-induced DNA damage [47]. The interaction of Cx43 with PRP19 and CDC5L suggests that Cx43 form part of the PRP19-CDC5L complex that plays a role in DDR [47]. Interestingly, the CDC5L subunit of the complex directly interacts with the master checkpoint kinase ATR [48, 49].The identification of several strong Cx43-binding factors involved in DNA repair suggests that Cx43 somehow affects genome stability by participating in or interfering with DDR.

Pull-down of Cx43 in nuclear extracts also revealed that Cx43 interacts with key proteins involved in DNA repair, such as Ku70 (XRCC6,) and the poly [ADP-ribose] polymerase 1 (PARP1), which are involved in DNA double-strand break repair (DSBR) and single-strand break repair (SSBR), respectively. However, DDX5, XRCC6 and PARP1 were more enriched in the Cx43 interactome of control cells (Fig. 3k and Fig. S5a), which contains significantly lower levels of Cx43, is mainly SUMOylated, and not found in the membranes (see Fig. 1c-d and 3i). These results suggest that Cx43 (and SUMOylated Cx43) favours certain interactions such as with LMNA and several others involved in DDR (Fig. 3m). Western blot and confocal microscopy assays confirmed the colocalization of Cx43 with LMNA and Ku70 and HMGB1 proteins, which were identified as Cx43-interactors by mass spectrometry (Fig. 3l-3m and S5b). Interestingly our results showed an increase in LMNA levels in Cx43-overexpressing cells (Fig. 3m and S5c).

DNA repair occurs within different nuclear compartments at the nuclear periphery [50] and the recruitment of the DNA damage sites to the nuclear lamin impairs DNA repair [50, 51]. Our results showed that an important fraction of nuclear Cx43 is bound to the chromatin (Fig. 3i) and additionally interacts with the chromatin, the nuclear lamin, several proteins involved in DDR and other complexes including HMGB1 [52], suggesting that Cx43 is involved in anchoring DNA repair complexes to the nuclear periphery, delaying or interfering with the pathways involved in DDR [50].

### Cx43 enhances the response to BRAF/MEK inhibitors treatments overcoming drug resistance

Since tumour cells under BRAF inhibitors treatments display increased sensitivity to DNA-damaging agents [53, 54] (Kong et al, 2007), we next sought to determine whether the upregulation of Cx43 in tumour cells harbouring the BRAF^V600E^ mutation further increases the DNA-damaging susceptibility under treatment with the BRAF/MEKi dabrafenib and trametinib, which are used in the clinic to treat BRAF-driven tumours [55, 56] (Fig. 4). Different BRAF^V600E^ tumour cells responded to 100 nM dabrafenib (D) and 1nM trametinib (T) for 48 h, as evidenced by a decrease in pRB and pERK (Fig. S6a). Increased levels of DNA damage, as measured by the comet assay, were detected in BRAF^V600E^-Cx43 tumour cells compared to control cells under treatment with 100 nM dabrafenib in combination with 1 nM trametinib for 72 h (Fig. 4a), consistent with DNA damage signalling pathways activation detected by 53BP1 nuclear foci formation and ATM activation (pATM) (Fig.4b). Moreover, Cx43 upregulation displayed the greatest reduction in the ability to form colonies of BRAF^V600E^ tumour cells after 7 days of BRAF/MEKi treatment (Fig. 4c). These results were recapitulated in different tumours cells harbouring the BRAF^V600E^ mutation (Fig. S6b-d). Cx43 decreased from 2.3 to 0.7 nM and 0.2 to 0.08 nM the half-maximal (50 %) inhibitory concentration (IC50) for dabrafenib and trametinib respectively (Fig. 4c and S6e). The decrease in colony formation in the presence of BRAF/MEKi was associated with an increase in cell death as detected by flow cytometry (Fig. 4d and S6f) and by an increase in cleaved caspase 3 (CC3) protein levels and CytC release (Fig. S6g), as well as an increase in cellular senescence (Fig. S6h-i).

**Figure 4.**
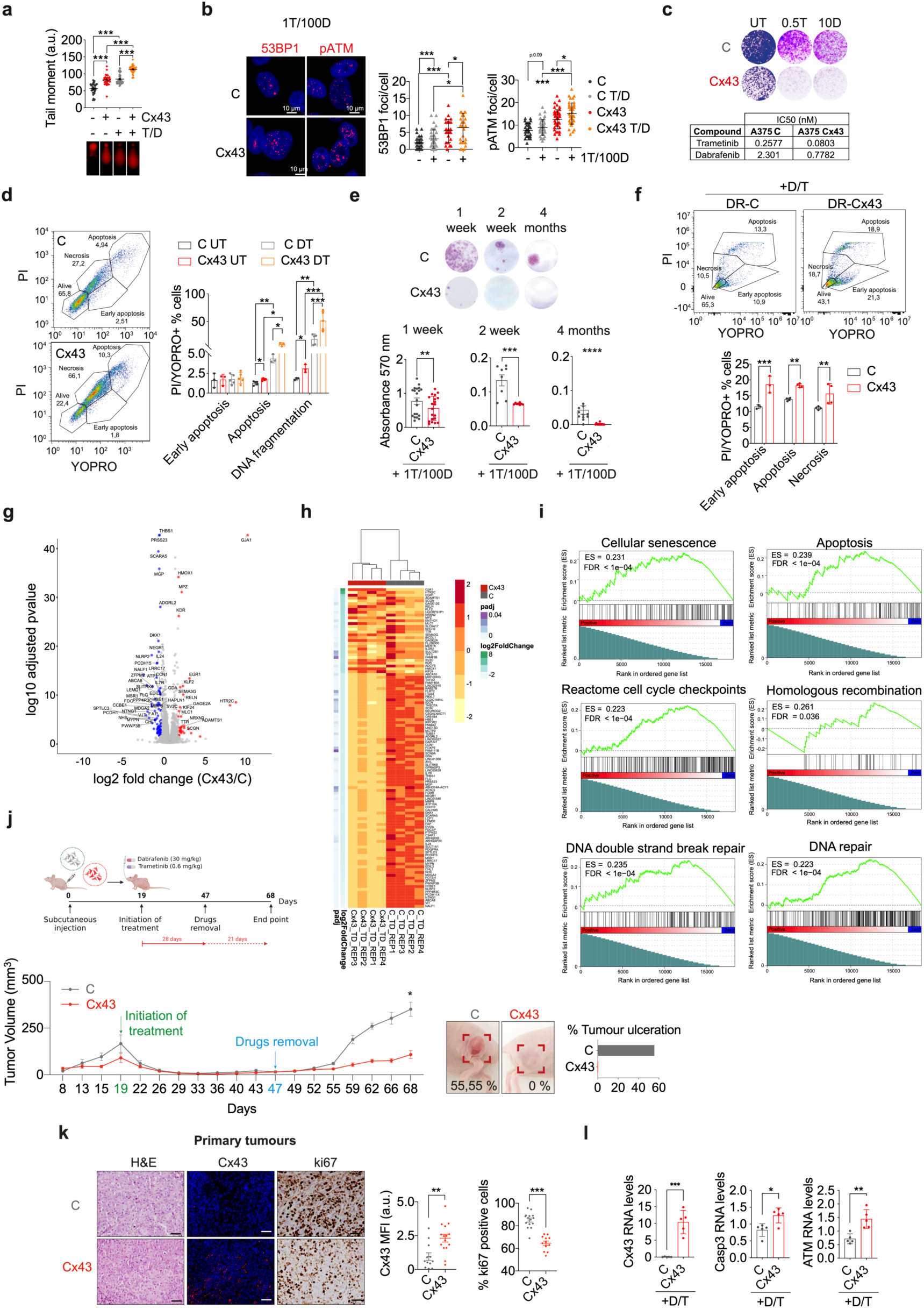
Cx43 enhances the response to BRAF/MEK inhibitors treatments and overcomes drug resistance. (a) Representative images from the comet assay analysis to evaluate DNA damage in control and Cx43 overexpressing cells. Cells were treated with 1 nM T and 100 nM D for 72 h. Tail moment was measured with the OpenComet plug-in from ImageJ software. The comets that were wrongly recognized by the program were removed manually. Data is representative of three independent experiments. **(b)** Representative nuclei images showing 53BP1 and pATM foci in C and Cx43 overexpressing A375 cells. An increment on the number of foci per cell is observed in Cx43 overexpressing cells in combination with BRAF/MEKi. n=3-6. **(c)** Colony formation analysis of A375 transfected with an C of Cx43 and treated with BRAF/MEKi. A significant cytotoxicity in the A375-Cx43 cell line (Dabrafenib IC50 = 0.7782 nM and Trametinib IC50 = 0.0803 nM) was detected compared to A375-C (dabrafenib IC50 = 2.301 and trametinib IC50 = 0.2577 nM). **(d)** Cell death measured by flow cytometry showing A375-EV and Cx43 cells undergoing early apoptosis, apoptosis and necrosis. Quantification of n=3-5 experiments is showed on the right. **(e)** Colony formation assays of A375-C and Cx43 cells after 1 week, 2 week and 4 months under 1 nM T and 100 nM D treatments. Data is representative of n=3. **(f)** Flow cytometry analysis showing DR cells transfected with Cx43 and its empty vector counterpart, undergoing early apoptosis, apoptosis and necrosis. (**g**) Volcano plot showing differentially regulated transcriptomics comparing control and Cx43 overexpressing A375 cells under BRAF/MEKi treatment (LFC ≥ 1 and ≤ -1; log10pvalue < 0.05). Signature proteins overexpressed in A375 C D/T are shown in blue and signature proteins overexpressed in A375 Cx43 D/T are shown in red. (h) Heatmap showing differentially expressed proteins (DEGs) comparing A375 C and Cx43 treated with BRAF/MEKi. DEGs were ranked based on their LFC. DEGs were included if adjusted p-value < 0.05 and |LFC| >=1. (i) GSEA showing positive enrichment of cellular senescence, apoptosis, cell cycle checkpoints, homologous recombination, DNA double-strand breaks and DNA repair signatures in A375-Cx43 D/T cells. (j) At the top, schematic representation of the workflow performed in the animal model. Middle panel, graph representing the tumour volume at the different stages of the experiment. Bottom, representative images and quantification of the percentage of ulcerated tumours in mice injected with A375 C compared to Cx43 overexpression. (k) Hematoxylin-eosin staining (H&E); Ki67 and Cx43 IF in primary tumours from mice injected with A375 C and Cx43 cells after treatment with BRAF/MEKi (TD). Cx43 levels were quantified as mean fluorescence intensity (MFI) and Ki67 as percentage of positive cells. (l) Cx43, CASP3 and ATM mRNA levels in tumour derived mice at endpoint, after 28 days of BRAF/MEKi treatment. All experiments were performed in at least three independent replicates. Data are expressed as mean ± SEM. Two-tailed Student’s t-test and one-way ANOVA were used to calculate significance as follows: *P < 0.05, **P < 0.01, ***P < 0.0001.

As expected, based on our previous results, when BRAF^V600E^-Cx43 cells were cultured in the presence of high concentrations of BRAF/MEKi (1T/100D to obtain resistant cells), we were unable to detect colonies after 2 weeks and 4-6 months in culture (Fig. 4e). To further investigate whether Cx43 plays a role in drug resistance to BRAF/MEKi treatments, BRAF^V600E^ mutant cells were treated with increasing concentrations of dabrafenib and trametinib for six months in monolayer culture, in order to obtain double resistant cells (DR) (Fig. S7a). The selected DR cells were resistant to very high concentrations of BRAF/MEKi (2 nM T/200 nM D and 5 nM T/500 nM D) detected by colony formation assays (Fig. S7a). When DR cells were transfected with Cx43, we observed increased sensitivity to cell death, as detected by flow cytometry (Fig. S7b-c). Our results showed that DR cells are slow-cycling cells (lower levels of pRB than control cells) with an epithelial to mesenchymal transition (EMT) phenotype that hardly express Cx43 (Fig. S7d). The restoration of Cx43 resulted in re-sensitisation of these DR cells to cell death and BRAF/MEKi treatment by increasing apoptosis and necrosis (Fig. 4f). These results demonstrate that Cx43 enhances BRAF/MEKi efficacy, overcomes resistance and re-sensitizes DR cells to cell death.

To confirm these results and to gain a more complete understanding of the genes and pathways affected by Cx43 in combination with BRAF/MEKi therapy, we performed RNA-Seq analysis to assess the gene signature after 24 h of combined treatments (Fig. 4g-i). We observed that the presence of Cx43 generated a differential footprint independent of the treatments, as shown in the volcano plot (Fig. 4g). This independent differential footprint suggests that Cx43 has the ability to regulate gene expression of different signalling pathways. Among the downregulated genes in BRAF^V600E^-Cx43, we identified several proteins involved in G1/S DNA damage checkpoints (e.g. HBE1, LCP2, THBS1, ZFPM2), cell cycle (e.g. CHL1, ARHGAP20, ARHGDIB) or MAP kinase tyrosine/serine/threonine phosphatase activity (DUSP6), among others (Fig. 4h). GSEA analysis confirmed our previous results, as we observed that the combination of Cx43 and BRAF/MEKi increased the representation of genes involved in cellular senescence, apoptosis, cell cycle checkpoints, homologous recombination, DDSB repair and DNA repair among other related pathways reinforcing the concept of specific synthetic lethalities (Fig. 4i).

To further confirm the activity of Cx43 in BRAF/MEKi responses, we designed a preclinical *in vivo* model. BRAF^V600E^ mutant control cells and Cx43-overexpressing cells were implanted subcutaneously in nude mice (Fig. 4j). After 19 days of injection, animals were treated with dabrafenib (210 mg/kg) and trametinib (4,2 mg/kg) by oral gavage for 28 days, showing a complete response (Fig. 4j). After 19 days of treatment resulting in a complete response, the drugs were removed, and the animals were monitored every 2 days for the following 20 days. The onset of relapse or tumour regrowth following treatment, suggested that BRAF^V600E^ mutant cells either exhibit a persistence of dormant tumour cells that resist treatments or develop resistance to the combined treatments by day 8 after drug removal (day 55 post-injection) (Fig. 4j). Notably, the presence of Cx43 resulted in a 70 % reduction (69.22) in tumour regrowth and relapse, indicating a significant protective effect (see Fig. 4j). Additionally, Cx43-xenografts exhibited no ulceration, a predictive marker for treatment response and prognosis [57] (Fig. 4j). IHC studies confirmed a decrease in Ki67 and an increase in ATM and CASP3 in BRAF^V600E^-Cx43 xenografts, 68 days after tumour cell injection and 21 days after drugs removal (Fig. 4k-l).

### Cx43-sEVs-based therapy in combination with BRAF/MEKi

Therapeutic protein and mRNA delivery using sEVs is enabling the development of highly effective therapies for various protein and gene therapeutics [58, 59]. These membrane-bound nanoparticles are relatively inert, non-immunogenic, biodegradable and biocompatible [60–62]. Based on current knowledge, we designed a proof-of-concept approach to restore Cx43 by introducing Cx43 into tumour cells and combining this treatment with BRAF/MEKi (Fig. 5). We ectopically expressed Cx43 in parental tumour cells (autologous treatment), which now produced sEVs enriched for Cx43 protein and mRNA transcripts (Fig. 5a). In vitro treatment of A375-BRAF^V600E^ mutant cells with Cx43-positive sEVs showed incorporation of Cx43 protein and mRNA into the target tumour cells as detected by western-blot and RT-qPCR (Fig. 5b) and correlated with a significant decrease in the ability to form colonies when cells were treated with Cx43-sEVs every 48 h over a period of 7 days (Fig. 5c). Cellular uptake of therapeutic sEVs was studied by labelling the nanovesicles with the fluorescent dye DilC and imaging by fluorescence microscopy (Fig. 5d and 5e). sEVs enriched with Cx43 from a different source (a non-tumoral cell line containing Cx43-enriched sEVs) reduced the capacity of tumour cells to form colonies in both A375-BRAF^V600E^ cells (Fig. 5f) and DR cells (Fig. 5g). These results showed that the delivery of mRNA and Cx43 protein using sEVs is an effective strategy and a potential therapeutic partner to increase efficacy and to overcome resistance to BRAF/MEKi.

**Figure 5.**
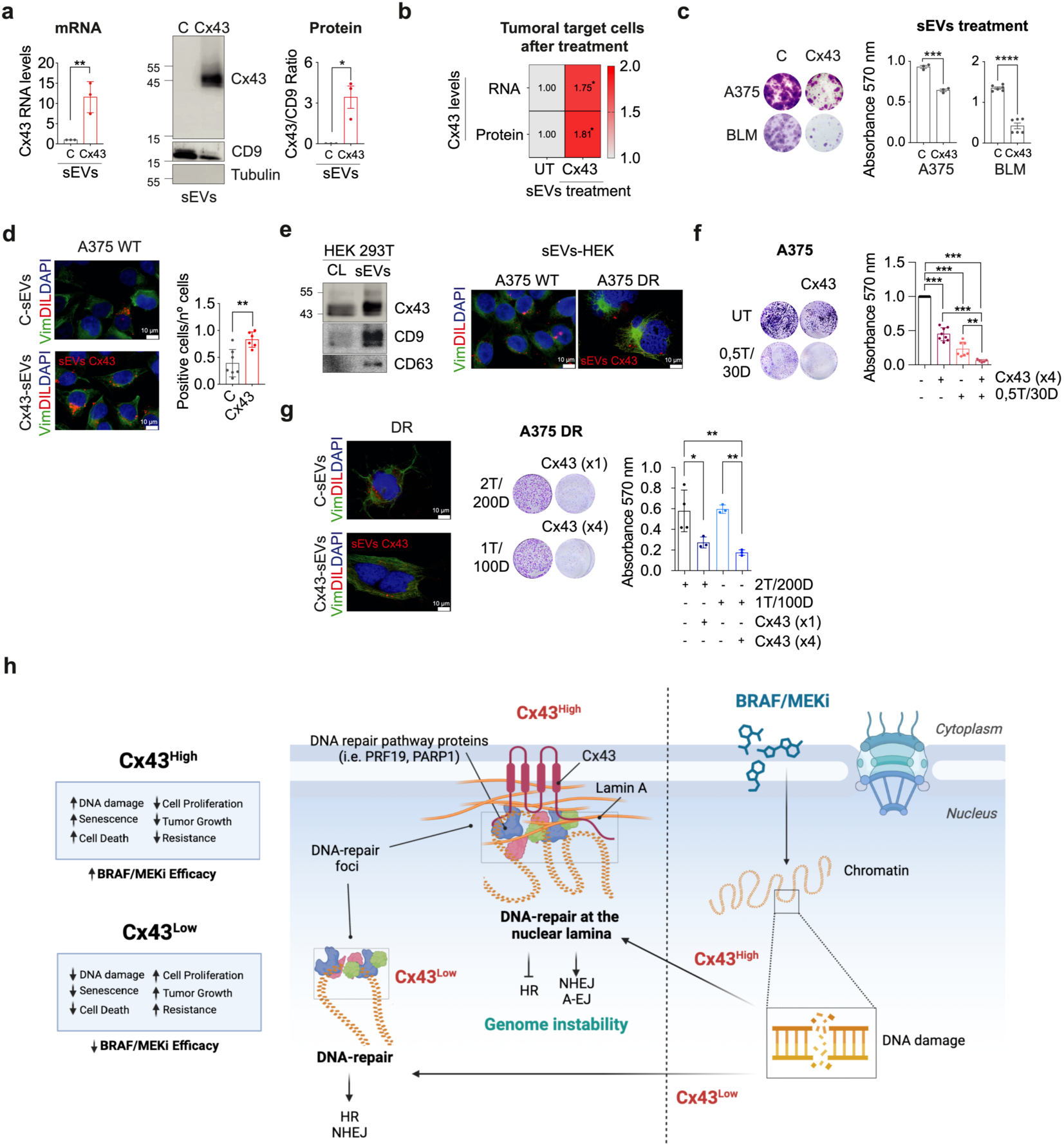
The presence of Cx43 in sEVs switches the role of these vesicles promoting the anti-tumour activity. (a) Cx43 mRNA and protein levels were analysed in sEVs derived from A375-C and Cx43 cells by RT-qPCR and western-blot, respectively. CD9 was used as sEVs marker and Tubulin was used as cell lysate loading control (n=3). **(b)** Heatmap showing Cx43 mRNA and protein levels in A375 untreated cells compared to A375 cells treated with Cx43 positive sEVs for 5 days. 2.22 x 10^12^ particles were used and the treatment was refreshed every 48 h. Representative data of three independent experiments is shown. **(c)** Colony formation assay of A375 and BLM tumour cell lines after 7 days treatment with Cx43-sEVs (2.23 x 10^12^ particles) or sEVs (2.36 x 10^12^ particles). n=3 independent experiments. **(d)** Vimentin (Vim, green) immunofluorescence was performed in A375 cells after 2 h treatment with Dil-labelled (red) sEVs to evaluate sEVs internalization. **(e)** Cx43 protein levels (on the left) were evaluated by western blot in cell lysates and sEVs derived from HEK293 cells; CD9 and CD63 were used as sEVs loading controls (n = 3). On the right, vimentin immunofluorescence (green) was performed in A375 and A375 DR cells after a 2 h treatment with Dil-labelled (red) HEK-derived sEVs. **(f)** Colony formation assay of A375 cells either untreated (UT), treated with (0.5 nM T/30 nM D) Cx43-sEVs (9.99 x 10^11^ particles) or with the combination of BRAF/MEKi and HEK-derived Cx43-sEVs for 7 days. Treatments were refreshed every 48 h. (g) On the left, sEVs internalization experiments showing vimentin immunofluorescence (green) in A375 DR cells after 2 h treatment with Dil-labelled (red) HEK-derived sEVs. On the rigth, colony formation assay of A375 DR cells, either treated with (2 nM T/200 nM D) and with the combination of BRAF/MEKi and HEK-derived Cx43-sEVs (9.99 x 10^11^ particles (x1) and 39.96 x 10^11^ particles (x4)) for 7 days. Treatments were refreshed every 48 h. **(h)** Proposed model of Cx43 based therapy mechanism in combination with BRAF/MEKi for promoting cell death and bypass drug resistance. In this study, we found that Cx43 interacts with DNA repair proteins in BRAF-mutated tumour cells, which can result in persistent DNA damage. The choice of the DNA repair pathway is influenced by how the DNA is organized in the nucleus, which is determined by Cx43 interactors and its recruitment to DNA damage sites. We have discovered a chemical synthetic lethality between BRAF/MEKi and Cx43, which when combined suppress these pathways (HR, NHEJ) and lead tumoral cells to apoptosis. All experiments were performed at least three independent times. Data is presented as mean ± SEM. Two-tailed Student’s t-test and one-way ANOVA were used to calculate the significance represented as followed: * P < 0.05, **P < 0.01, ***P < 0.0001.

## Discussion

In this work, we have shown that Cx43 reduces tumour growth in BRAF-mutated tumours, enhances BRAF/MEKi sensitivity and overcomes mechanisms associated with drug evasion and resistance by increasing genome instability and enhancing tumour cell lethality in both sensitive and resistant cells (Fig. 5h).

Melanoma driven by BRAF^V600E^ mutation is a widely used model to study drug adaptation and resistance in cancer models. Both BRAF driver mutations and BRAF/MEKi induce cellular senescence and DNA damage [53, 63, 64]. Our results reveal that in a malignant context with sustained DNA damage, such as under BRAF and NRAS mutations, Cx43 enhances cell senescence, and predominantly increases susceptibility to cell death under BRAF/MEKi treatment. Our results were consistent across all cell lines analysed, suggesting that the effect of Cx43 is neither drug- nor cell-specific (Fig. 3f and S6). In particular, we show that Cx43 is an important therapeutic target in these tumours and that not only dramatically enhances the efficacy of BRAF/MEKi in BRAF-mutant malignancies with varying sensitivity to these agents, but also cooperates with MEK inhibitors in NRAS-mutant tumours. Further, cells that rapidly escape drug by non-genetic mechanisms are prone to DNA damage [65]. Thus, we have shown that in this setting, Cx43 increases the susceptibility to cell death of both sensitive (naive) and resistant cells, alone and in combination with BRAF/MEKi. Increasing the vulnerability of tumour cells in the context of therapy will allow for the development of new strategies that lead to the elimination of minimal residual disease, which is responsible for driving therapy relapse. Cx43 modulation therefore represents an excellent opportunity to be exploited in order to improve the clinical benefit of targeted therapies that induce DNA damage in cancer, such as BRAF/MEKi.

Although we have previously found that Cx43 interacts with several nuclear and perinuclear proteins, including factors involved in chromatin organization or proteins of the nuclear envelope and lamins [46], in this study, we discovered that Cx43, in the tumour cells, interacts with various DNA repair proteins resulting in persistent DNA damage, which in turn results in synthetic lethality under BRAF/MEKi treatment. The choice of the DNA repair pathway has been shown to be controlled by the spatial organization of the DNA in the nucleus [50, 51]. The chromatin structure and mobility capacity at the inner nuclear lamin leads to a slower DDR and restriction of the HR, allowing preferentially NHEJ and A-EJ [50], which are considered highly mutagenic pathways, instead of the error-free HR pathway. Our results suggest that Cx43 acts as a signaling hub with many interaction partners, including the nuclear lamin, proteins involved in various DNA repair pathways (i.e. PRPF19, PARP1 or XRCC6) and several histones such as histone H2B type 3-B or histone H2A type 1, involved in the recruitment of repair proteins to sites of DNA damage. Interestingly, we also found that Cx43 binds to the chromatin and interacts with a major component of the mammalian chromatin, HMGB1, that besides participating in DNA repair and genome stability, has also been involved in modulating the balance between senescence and cell death by apoptosis in response to DNA damaging agents such as doxorubicin or camptothecin [66].

Cx43 interacts with different transcription factors such as the CCAA/enhancer-binding protein zeta (CEBPZ) or the elongation factor 1-gamma (EEF1G) and the RNA-seq shows that Cx43 affects a broad transcriptional network of different pathways including genes involved in DDR and DNA repair. Our study identifies Cx43 as a potential factor involved in the spatial organization of lamin-associated chromatin domains related to the regulation of DDR and genome instability. Given that Cx43-interactors include topoisomerase 1 and several splicing factors (i.e. FUS, PRP19 or several hnRNAPs) that cooperate as gatekeepers of genome stability [67], further studies are needed to determine the potential role of Cx43 in mRNA splicing and transcription-coupled DNA repair.

DNA damage can induce a tumor-suppressive response termed cellular senescence, which harbors persistent nuclear foci containing DDR proteins. Our results show that upregulation of Cx43 significantly enhances cellular senescence in BRAF- and NRAS-mutant tumours. However, our data also reveal that Cx43 provides a unique opportunity for cancer therapy by inducing persistent DNA damage and synthetic lethality bypassing senescence under BRAF/MEKi treatment. The mechanism by which Cx43 induces senescence can be attributed to the anchor of DNA repair foci to the nuclear lamin, thus increasing DNA damage and persistent nuclear foci (by interfering with DNA repair pathways), or as result of changes in gene expression, or a combination of both. Regardless, our results demonstrate that the combined suppression of BRAF/MEK and Cx43 restoration effectively capitalizes on this vulnerability.

Cx43 is known to change gene expression [68] and has been associated with reduced proliferation in different tumour types [25, 69, 70]. However, the molecular mechanism involved in its tumour suppressor role and in the response to therapy remains unknown [71]. Our findings mechanistically dissect its nuclear role and reveal Cx43 as a novel player in the regulation of the DDR. Another conclusion from our study is that Cx43 bounds directly to chromatin and changes the landscape of gene expression. Although the role of Cx43 in modulating chromatin structure or acting as a transcription factor has not been reported, previous reports are aligned with these functions, since it regulates the expression of specific genes and it was reported to interact with subunits of RNA-polymerase II [68]. Overall, these results expand our understanding of how Cx43 exerts its anti-tumour activity and affects the targeted therapy responses. However, additional work is needed to further determine the role of Cx43 in gene transcription and DNA repair.

Endogenous nanoparticles termed sEVs are considered the most promising next-generation nano-delivery tools and Cx43 has been shown to be required for sEVs cargo release into recipient cells [72, 73]. Interestingly, when Cx43 was overexpressed in BRAF-mutant tumour cells using a vector, the sEVs secreted by the tumour cells were enriched for both Cx43 mRNA and protein. The presence of Cx43 in the secreted sEVs changes the small RNA and protein content of these vesicles and their role (unpublished results from our group). In this study, the anti-tumour effect of the Cx43-enriched sEVs was exploited as a new therapeutic approach, as sEVs have many advantages such as small size, good biocompatibility, low toxicity, and low immunogenicity making them ideal for delivering mRNA molecules and membrane proteins (e.g. NCT05156229)[58]. The data demonstrate that delivering Cx43 to both sensitive and resistant cells decreases their ability to form colonies and notably enhances the effectiveness of the inhibitors. This offers a proof-of-concept in relevant preclinical models. In summary, our results demonstrate that Cx43 inhibits the growth of BRAF-mutant tumours and NRAS-driven melanoma, a cancer type for which effective therapies are lacking. Furthermore, our study reveals that Cx43 significantly enhances the efficacy of and sensitizes tumour cells to cell death induced by BRAF/MEKi, regardless of their sensitivity or resistance to these agents. Additionally, we present a novel strategy to restore Cx43 in tumour cells using sEVs as a drug delivery system. We have also elucidated the mechanisms underlying Cx43 function and identified a novel player in DDR. Collectively, these findings present a promising and translatable strategy for enhancing treatments for RAS/RAF pathway-driven tumours.

## Acknowledgements

This work was supported in part through funding from a grant from Ministry of Science, Innovation and Universities (MICIU/AEI/10.13039/501100011033): PID2022-137027OB-I00 ERDF/EU and a grant from the Joint Transnational Call for Proposals for “European Innovative Research & Technological Development Projects in Nanomedicine” EURONANOMED III (AC21_2/00026) (to MDM); a grant from Xunta de Galicia (IN607B2020/12) (to MDM); a grant from HORIZON-MSCA-2023-SE-01 (101183034) and FET-OPEN-Future and Emerging Technologies (to MDM) and from EU HORIZON-CSA 101079489. TWINFLAG (to MDM). AV-V, PC-F and MV-E were funded with post-doctoral fellowship (IN606B 2022/002; INB606B 2017/014 and IN606C 2021/006; IN606B-2019/004) from Xunta de Galicia. AG-C was funded with a predoctoral fellowship (FIS20/00310) from ISCIII. AC-F was funded with a predoctoral fellowship (IN606A 2022/038) from Xunta de Galicia. We thank members of the CellCOM group for helpful technical suggestions, Biobank A Coruña for collecting tissue samples, Arantxa Tabernero (INCYL, University of Salamanca) for kindly providing the human Cx43 plasmid used in this study, the group of Dr. María Pardo (Obesidomic Group, IDIS, Santiago de Compostela, Spain) for advise us in the first step of sEVs isolation and characterization, the group of Dr. José Luis Labandeira (Cell and Molecular Neurobiology of Parkisońs disease, CiMUS, Santiago de Compostela, Spain) for help with *ex vivo* assays, the Dr. Julián Yañez and Dr. María Luz Díaz (NEUROVER Group, University of Coruña, Spain), Dr. Ángel Fernández (Department of Cellular Pathology, Hospital La Reina, Ponferrada, Spain), IDIS facility (María Otero and Dr. García Caballero, Santiago de Compostela, Spain) and María C. Vázquez (Biobank A Coruña, Spain) for the knowledge in the area of anatomical pathology and tissue histology. The Genomics Unit CNIC (Dr. Alberto Benguría and team) for their help and support. We would also like to acknowledge Dr. Silvia Penuela (University of Western Ontario, Canada) and Dr. Edward Leithe (Oslo University Hospital HE, Norway) for providing vectors and antibodies. Although these were not ultimately included in the results of this manuscript, their valuable input significantly contributed to our research.

## Author contributions

AV-V and AG-C together with PC-F, VA, AC-F, MV-E and MR-CM designed and performed experiments, analyzed data, and prepared figures. TC, MQ, JRC and EF provided input, clinical advice and facilitated access to BRAF/MEK inhibitors (TAFINLAR and Mekinist) for *in vitro* and *in vivo* experiments. SB-L performed and advised with proteomics data. DS, AV and JS-L contributed with support and helpful for *in vivo* models. GS and CF-L provided input and supported RNA-seq assays and performed data analysis. MGB and PH contributed with the knowledge in DNA repair mechanism. BS-L participated with critical input and in the design of *in vitro* and in vivo models. CA provided critical input and performed together with VA the experiments and analysis related with DNA damage and binding to chromatin. MDM conceived, directed and supervised the study. MDM wrote the manuscript with the support of EF. AV-V, AG-C and with the input from all co-authors. All authors reviewed the manuscript.

## Data availability

The data that support the findings of this study are available from the corresponding author upon reasonable request.

## Methods

### Cell lines and culture conditions

All cell lines used in this study are listed in TableS1 Cell lines were cultured under standard conditions in Dulbecco’s Modified Eagle Medium (DMEM) (Gibco) medium supplemented with 10 % Fetal Bovine Serum (FBS) (Life Technologies Gibco) and 100 U/mL penicillin (Life Technologies) and 100 µg/mL streptomycin (Life Technologies), unless otherwise indicated. HT-29 human colorectal adenocarcinoma cells were grown in McCoy’S 5A medium (Sigma-Aldrich) supplemented with 10 % FBS and 100 U/mL penicillin and 100 µg/mL streptomycin. All cell lines were regularly controlled to be mycoplasma-free and previously sequenced to verify authenticity and avoid cross-contamination.

### Generation of double resistant cell line

Double resistant cells were generated after exposure of very low confluence of parental A375 cells to increasing concentrations of the BRAF inhibitor dabrafenib (Tafinlar® 75 mg, Novartis) and the MEK inhibitor trametinib (Mekinist® 2 mg, Novartis) (5 nM to 200 nM dabrafenib plus 0.1 nM to 2 nM trametinib) until cells resumed growth. Double resistant cells were finally maintained using 200 nM dabrafenib and 2 nM trametinib for a minimum of 3 months before considering them resistant. After acquiring the resistant phenotype, they continued to be cultured under BRAF/MEKi.

### Cell transfection

A375, SK-Mel-28, SK-Mel-103, SK-Mel-147, BLM, MDA-MB-231 and HT-29 cell lines were transfected with a plasmid to overexpress Cx43 kindly provided by Dr. Arantxa Tabernero (Institute of Neuroscience of Castilla y León, University of Salamanca, Spain) as previously described [19]. Cells were subsequently selected with 0.5 – 1 µg/mL puromycin dihydrochloride (Tocris).

### Animal studies

All animals were maintained under specific pathogen-free conditions and handled in accordance with the Animal Welfare guidelines and were performed in accordance with protocols approved by The University of Santiago de Compostela Bioethics Committee in compliance with Principles of Laboratory Animal Care of national laws (license number 15002/2019/009). For tumour xenograft experiments, cells were resuspended using 30 % Matrigel in DMEM 10% FBS, and 1 x 10^6^ of A375-C or A375-Cx43 were subcutaneously injected into the right flank of 6 to 8-week-old female nude BALB/c mice (NMRI-Foxn1nu/nu, Janvier Labs). Tumours were allowed to establish, sizes (average 50 mm^3^) were matched and then mice were randomly allocated to groups of 4-5 animals. Treatment was administered by orogastric gavage with 30 mg/kg Tafinlar (Dabrafenib, Novartis) and 0.6 mg/kg Mekinist (Trametinib, Novartis). Drugs were dissolved in 0.5 % methylcellulose and 0.2 % Tween 80. Drugs were administered daily, 7 days a week. Tumour volume was determined twice a week (V = (W^2^ x L)/2)). At the endpoint of each group of animals, tumours were weigthed and divided in two: half tumour was formalin-fixed and paraffin-embedded using standard protocols and the other half was immediately frozen to RNA extraction.

### RT-qPCR analysis

Total RNA was collected and purified using TRIzol (Sigma) according to the manufacturer’s instruction. 200 μl of chloroform were added to all the samples and were subjected to vigorous agitation for 15 s before being incubated at room temperature (RT) for 3 min. To guarantee the absence of DNA in the samples, the RNA was treated with DNase (Invitrogen’s RNase-free DNase). The SuperScript® VILO^TM^ cDNA Synthesis Kit was utilized according to the manufacturer’s guidelines (Invitrogen) to synthesize cDNA, with a total of 1 μg of total RNA per reaction. RT–qPCR was performed using LightCycler 480 (Roche). RNA expression was normalized to the expression of HPRT1 and actin (RT–qPCR primers are listed in Table S2).

### Immunoblot analysis

Cells were harvested and physically lysed through 30-gauge syringes in lysis buffer (150 mM NaCl, 50 mM Tris-HCL (pH 7.5), 5 mM EDTA (pH 8), 0.5 % (v/v) Nonidet P-40, 0.1 % (w/v) Sodium Dodecyl Sulfate (SDS) and 0.5 % (v/v) N-Lauroylsarcosine) supplemented with 5 µg/mL protease inhibitors cocktail and 1 mM PMSF. Total protein content was determined with BCA Protein Assay Kit (Thermo Scientific) according to the manufacturer’s protocol. Equal amounts of protein were separated in 10-15 % SDS-PAGE under reducing conditions and transferred onto a polyvinylidene fluoride (PVDF) membrane (Inmobilon-P, Millipore) using Mini Trans-Blot Cell System (Bio-Rad Laboratories). Transfer was performed at 120V for 1 h at 4°C. Membranes were blocked with 5 % milk in tris-buffered saline with 0.05 % Tween-20 (TBS-T) for 30 min at RT. Primary and secondary antibodies (Table S3) were diluted in 5 % milk TBS-T incubated overnight at 4°C or 1 h at RT. Protein bands were detected by the ECL system (PierceTM ECL Western Blotting Substrate, Thermo Scientific™) and visualized with Amersham Imager 600 (GE Healthcare).

### Immunofluorescence and immunohistochemistry analysis

Cell immunofluorescence was performed as previously described [33]. Briefly, cells were seeded onto coverslips, fixed for 15 min at RT with 4 % paraformaldehyde (PFA), washed with Phosphate-buffered saline (PBS), pH ∼ 7.4 and blocked for 30 min at RT with blocking buffer (1 % BSA in PBS). Fixed and membrane-permeabilized cells were incubated with listed primary antibodies for O/N at 4°C, and with fluorescent-secondary antibodies for another 1 h at RT, in the dark. Nuclei were stained with 4’,6-diamidino-2-phenylindole dihydrochloride (DAPI; Sigma-Aldrich) for 5 min at RT in the dark. Images were either obtained in an Olympus BX61 microscope using a DP71 digital camera (Olympus) or in a confocal microscope A1R (Nikon) coupled to an inverted microscope model Eclipse Ti E (Nikon). After data collection, ImageJ and FIJI software were used for further analysis. For High throughput microscopy, cells were seeded at 10,000 cells per well in clear bottom, black 96-well plates (Greiner, 655090). 24 h later cells were treated with 3 mM Hydroxyurea (Sigma, H8627), 1μM Camptothecin (Sigma, C9911), 25 μg/mL Bleomycin (Stratech, S1214), 25 μg/mL Aphidicolin (Sigma, A4487) and 25 μM Mitomycin-C (Duchefa Biochemie, M0133) for 2 h. To label S-phase cells, 20 μM EdU was added to the media for 20 min. Fixation was performed for 20 min with 2 % formaldehyde and cells are permeabilized in 0.2 % Triton in PBS. If pre-extraction is required, cold 0.5 % Triton X-100 CSK buffer (10 mM PIPES pH 7, 100 mM NaCl, 300 mM sucrose, 3 mM MgCl_2_) with protease inhibitors is added for 5 min at 4 °C before fixation. In this case, permeabilization is not required. For EdU detection, Alexa-647 was covalently linked to EdU using Click-iT® EdU Imaging Kit (Thermo Fisher, B10184). Samples were blocked with BSA and primary and secondary antibody incubation are performed according to the manufacturer conditions as follow: anti-Connexin43 1:1000 (Merk, C6219), anti-γH2AX 1:400 (Millipore, 05-636), anti-Rabbit IgG Alexa Fluor 488 1:1000 (Invitrogen, 1910751), anti-Mouse IgG Alexa Fluor 546 1:1000 (Invitrogen, A11030). DAPI 1:400 (Thermo Fisher, 62248) was added together with the secondary antibodies, cells were washed with PBS-0.1 % Tween20, and left in PBS until imaging. Images were taken and analysed with ScanR High Content Screening Microscopy (Olympus). High throughput microscopy images were analysed with ScanR analysis software. Data were visualised and statistically analysed in Tableau and GraphPrism. Data shown in this work corresponds with the signal detected in the nuclear area, determined by DAPI intensity. For immunohistochemistry analysis, 10 µm paraffin embedded human tissue slides were stained with OptiView DAB IHC Detection kit (Roche). The slides were permeabilized with 0.1 % Tween20 in PBS. Then, we used the OptiView Peroxidase Inhibitor 3.0 %. The slides were washed three times with PBS. Next, it was incubated with primary antibodies for 1 h at RT or O/N at 4 °C. The slides were washed three times with PBS followed by an incubation with the secondary antibody OptiView HQ Universal Linker for 10 min at RT. The slides were washed three times with PBS and then we added the OptiView HRP Multimer for 8 min. After three washes with PBS, the slides were incubated with 0.2 % OptiView DAB in OptiView H2O2 for less than 1 min. Finally, the slides were washed with H_2_O and stained with hematoxylin/eosin (H&E). The antibodies used in this procedure are listed in TableS3.

For single cell confocal microscopy, after keeping cells growing to allow Cx43 overexpression at least 10 days, MCF7 cells were seeded at 10,000 cells per well in clear bottom, black 96-well plates (Greiner, 655090). 24 h later cells were treated with 3mM hydroxyurea (Sigma, H8627), 1 μM Camptothecin (Sigma, C9911), 25 μg/mL bleomycin (Stratech, S1214), 25 μg/mL aphidicolin (Sigma, A4487) and 25μM mitomycin-C (Duchefa Biochemie, M0133) for 2 h. To label S-phase cells, 20 μM EdU was added to the media for 20 min. Fixation was performed for 20 min with 2 % formaldehyde and cells are permeabilised in 0.2 % Triton in PBS. If pre-extraction is required, cold 0.5 % Triton X-100 CSK buffer (10 mM PIPES pH7, 100 mM NaCl, 300 mM sucrose, 3 mM MgCl_2_) with protease inhibitors is added for 5 min at 4 °C before fixation. In this case, permeabilization is not required. For EdU detection, Alexa-647 was covalently linked to EdU using Click-iT® EdU Imaging Kit (Thermo Fisher, B10184). Samples were blocked with BSA and primary and secondary antibody incubation are performed according to the manufacturer conditions as follow: anti-Connexin43 1:1000 (Merk, C6219), anti-γH2AX 1:400 (Millipore, 05-636), anti-Rabbit IgG Alexa Fluor 488 1:1000 (Invitrogen, 1910751), anti-Mouse IgG Alexa Fluor 546 1:1000 (Invitrogen, A11030). DAPI 1:400 (Thermo Fisher, 62248) was added together with the secondary antibodies, cells were washed with PBS-0.1 % Tween20, and left in PBS until imaging. Images were taken and analysed with ScanR High Content Screening Microscopy (Olympus).

### High throughput microscopy data

High throughput microscopy images were analysed with ScanR analysis software. Data were visualised and statistically analysed in Tableau and GraphPrism. Data shown in this work correspond with the signal detected in the nuclear area, determined by DAPI intensity.

### Flow cytometry analysis

Flow cytometry analysis data was collected on a FACScalibur^TM^ flow cytometer (Becton Dickinson) using a CellQuest^TM^ Pro software and the BD FACSCanto™ II system (Becton Dickinson) using a BD FACSDiva software and analyzed by FCS Express 6 Flow software (De Novo Software). For apoptosis assays, cells were harvested and resuspended in 300 µl of flow cytometry (FC) buffer after 3 PBS washes. Cells were labeled with double staining PI/YO-PRO-1 at 2 µg/mL and 150 nM, respectively. PI-/YO-PRO-1-cells were considered alive and apoptotic stages were considered based on YOPRO or PI incorporation levels. Alive cells were gated based on forward scatter (FSC) and side scatter (SSC) parameters to discriminate cell debris. Singlet cells were sorted from aggregates on the basis of FSC-H and FSC-A.

### Senescence-associated β-galactosidase activity

SA-βGal activity was assessed by flow cytometry with the fluorogenic β-galactosidase substrate di-β-D-galactopyranoside (FDG; Invitrogen, Thermo Fisher Scientific) or with the Senescence Cells Histochemical Staining Kit (Sigma-Aldrich), as previously described [19].

### Click-iT assay

Cell proliferation by tracking new DNA synthesis was performed with Alexa Fluor 647™ Click-iT® Assays kit (Thermo Scientific™) following manufacturer’s instructions. Cells were fixed and incubated with the Alexa 647™ picolyl azide. EdU incorporation was measured by flow cytometry with a FACScalibur^TM^ flow cytometer (Becton Dickinson) with the CellQuest^TM^ Pro software and analyzed with the FlowJo™ Software.

### Clonogenic assay

Colony formation assays were performed by seeding 5 ·10^3^ cells per well onto 6-12 well plates and grown for 7-15 days. For double resistant cells, colony formation assay was performed treating the cells with dabrafenib (Novartis) and trametinib (Novartis) from 1 week to 4 months, and then, the drug was removed for one week. The treatments were refreshed every 48 h. Cells were fixed with 2 % PFA for 15 min and stained with 0.05 % crystal violet. The stained cells were then rinsed with distilled water and left to dry O/N at RT. Colony images were captured using the Epson Perfection V800 photo scanner. Quantification was performed by diluting the crystal violet staining with 30 % acetic acid for 15 min at RT. Quantification was performed by measuring the absorbance at 570 nm using the NanoQuant microplate reader Infinite M200 (TECAN).

### Adhesion assay

Culture plates were pre-treated with Collagen Type I solution (Sigma-Aldrich) O/N at 4 °C and blocked with 1 % BSA at 37 °C for 1 h. Then, 2 · 10^5^ cells were plated at 37 °C for 2 h. Cells were fixed with 2 % paraformaldehyde and stained with 0.1 % crystal violet (Sigma-Aldrich). Colonies were quantified in a NanoQuant Infinite M200 (TECAN) microplate reader.

### Cell Migration assay

Cell migration capacity was measured using a Transwell^®^ Permeable 6.5 mm insert, 24 well plate and 8.0 µm polycarbonate membrane (Corning®). Polycarbonate membrane was pretreated with a Collagen Type I solution O/N at 4 °C for optimal cell attachment. 2 · 10^4^ cells were plated in the upper chamber with a porous membrane and incubated O/N at 37 °C. A chemoattractant, DMEM 10 % FBS is added into the bottom chamber to form a chemotactic gradient. Cells sense the chemotactic gradient and migrate through the pores of the transwell membrane. Migrated cells were stained with 0.1 % crystal violet (Sigma-Aldrich) and measured in an Infinite M200 (TECAN) microplate reader.

### Scrape loading/dye transfer (SL/DT) assays

Melanoma cells were seeded onto 6 or 12 wells plates and cultured until 80 – 100 % of confluence. Lucifer Yellow (LY) CH dilithium salt (1 mg/mL) (Cell Projects Ltd© Kent) in PBS was added to the cells and parallel cuts were performed with a surgical blade (Swann-Mortonscalpel®). After 3 - 5 min at 37 °C, cells were washed with PBS and fixed with 2 % PFA in PBS. LY becomes incorporated by the membrane-damaged cells along the cut and transferred into adjacent cells connected by functional gap junction channels. Images were captured in an inverted Nikon Eclipse Ti fluorescent microscope (Nikon). The score was calculated as the ratio of the number of non-damage LY positive cells at the edge and the number of the LY positive cells outside the edge.

### Dye Coupling Microinjection

Cells were seeded on 12-wells plates and cultured until 80-100 % of confluence. Glass micropipettes were backfilled with 133 mM of 5- or 6-carboxyfluorescein with a negative charge (-2) (Sigma-Aldrich) diluted in potassium acetate 0.1 M (Sigma-Aldrich). One cell was injected, and hyperpolarizing pulses were applied with an intensity of 0.25-1 nA. Cells were excited with a 488 nm laser in a confocal microscopy and the image was registered in Lasershap MRC-1024 (BioRad). The fluorescence emitted was measured at 640/40 nm and 540/40 nm with two photodetectors, and the time course of the passage of dye from the injected cell to the adjacent cells was evaluated.

### Determination of cellular ROS

Cellular superoxide content was measured using the DCFDA/H2DCFDA - Cellular ROS Assay Kit (abcam), following manufacturer’s instructions. Fluorescence-activated detection was carried out using Infinite M NANO + plate reader (TECAN) fluorimeter. Data is expressed as intensity of fluorescence.

### Hemichannel activity assay

ATP release levels were used to determine hemichannel activity. ATP levels were measured in triplicates in three independent experiments using the Adenosine 5′-triphosphate Bioluminescent Assay Kit (Sigma) according to the manufacturer’s instructions. Samples were measured using an Infinite 200 PRO (TECAN) luminometer.

### Protein immunoprecipitation

Cell lysates were cross-linked by adding 37 % formaldehyde and glycine and prepared in IP-lysis buffer (150 mM NaCl, 50 mM Tris-HCL (pH 7.5), 5 mM EDTA (pH 8), 0.5 % (v/v) Nonidet P-40, 0.1 % (w/v) Sodium Dodecyl Sulfate (SDS) and 0.5 % (v/v) N-Lauroylsarcosine) supplemented with proteinase/phosphatase inhibitors. IP was performed with 5 ·10^6^ cells by incubation with magnetic beads (BioMag®Plus Amine Protein Coupling Kit, BioTrend) previously conjugated with the antibodies indicated in Table S3. The preclearing process was performed to avoid unspecificities according to the manufacturer’s instructions. Cell lysate and antibody-conjugated magnetic beads were incubated O/N at 4 °C, followed by several washes in IP-lysis buffer. Complexes were eluted in Laemmli buffer and denatured at 75 °C, for 30 min before analysis.

### sEVs isolation, characterization, and *in vitro* delivery assays

A375-C, A375-Cx43 and HEK293T cells were washed 3 times with PBS, cultured for 48 h in DMEM without FBS (Gibco) and supernatants were collected for sEVs isolation. Cells, debris and large vesicles were first removed by serial spinning at 2000 *g* for 10 min and them filtered using a syringe filter with a pore size of 0.22 μm followed by ultracentrifugation in a Hitachi CP100NX ultracentrifuge at 100,000 *g* for 90 min. For experimental *in vitro* assays, the resulting sEVs pellet was resuspended in complete culture media and 8.3 x 10^12^ particles of sEVs approximately were added to receptor cells every 2 days. For the NTA Nanosight NS300 analysis and sEVs protein/RNA content determination, a second ultracentrifugation with PBS was performed, and the sEVs pellet was resuspended in PBS. For sEVs tracking, sEVs pellet was stained with 1 µM Dil (Invitrogen) for 1 h at 37 °C and visualized in an Olympus BX61 fluorescent microscope. For transmission electron microscopy (TEM) visualization, purified sEVs were fixed in 2 % PFA and deposited onto formvar-carbon coated grids, which were incubated with 0,1 M glycine, 1 % BSA and fixed with 1 % glutaraldehyde. Grids were stained in uranyl-oxalate at pH 7 and contrasted in a mixture of 4% uranyl acetate (Polysciences) and 2 % methyl cellulose (Sigma-Aldrich). sEVs were visualized in a JEM 1010 transmission electron microscope (JEOL).

### Proteomics and Mass Spectrometry Analysis

Cx43 IP of A375-C and Cx43 nuclear fraction (NUC) (n=4) were collected in Laemmli buffer for protein isolation. 100 µg from all conditions were simultaneously concentrated by 10 % SDS-PAGE and visualized by Sypro-Ruby fluorescent staining (Lonza). Gel slices were excised and, previous concentration, subjected to in-gel manual trypsin digestion following the protocol defined by Shevchenko [74] with minor modifications. Peptides were extracted by performing three 20-min incubation in 40 μL of 60 % ACN (Sigma-Aldrich) dissolved in 0.5 % formic acid (HCOOH) (Sigma-Aldrich). For mass spectrometry analysis (MS), peptides were analyzed by LC-MS/MS using a micro liquid chromatography system (Eksigent Technologies nanoLC 400, SCIEX) coupled to a high-speed Triple TOF 6600 mass spectrometer (SCIEX) with a micro flow source. Peptides were loaded onto a reverse phase column (Chrom XP C18 150 × 0.30 mm,3 mm particle size and 120 Å pore size; Eksigent, SCIEX), separated and MS data acquired using Analyst TF 1.7.1 software (SCIEX).

### Mass Spectrometry Data Processing and Analysis

For identification, data was processed using ProteinPilot^TM^ 5.01 software (SCIEX), with the algorithm Paragon^TM^ for database search and Progroup^TM^ for data grouping. Data was searched using a Human specific Uniprot database (https://www.uniprot.org/uniprotkb?query=human). False discovery rate (FDR) was performed using a non-lineal fitting method displaying only those results that reported a 1% Global false discovery rate (FDR) or better (shilov IV and Tang WH). Protein Quantification was performed using a Spectral count method. The search parameters of Protein Pilot are predefined in the software[75]. MS/MS were normalized between samples using Scaffold 5.0 by the sum of the unweighted spectral counts for each sample method in order to determine a sample specific scaling factor and then this was applied to all proteins in all the samples. Therefore Scaffold (version Scaffold_5.0.0, Proteome Software Inc., Portland, OR) was used to validate MS/MS based peptide and protein identifications. Peptide identifications were accepted if they could be established at greater than 95% probability by the Percolator posterior error probability calculation[76]. Protein identifications were accepted if they could be established at greater than 99% probability and contained at least 1 identified peptide. Protein probabilities were assigned by the Protein Prophet algorithm [77]. Proteins that contained similar peptides and could not be differentiated based on MS/MS analysis alone were grouped to satisfy the principles of parsimony. Proteins sharing significant peptide evidence were grouped into clusters. All calculations were performed using GraphPad Prism 7.0. Enrichment analyses were performed by STRING software version 11.0b (Gene Ontology (GO), KEGG Orthology and Reactome database).

### Gene expression analysis by RNA-Seq and computational data analysis

RNA was isolated from A375-C and A375-Cx43 untreated cell lines and under BRAF/MEKi (dabrafenib and trametinib) treatment (n=4, each condition) using the Kit RNA isolation: NucleoSpin RNA (Macherey-Nagel). Quality and integrity of each RNA sample was checked using a Bioanalyzer 2100 instrument (Agilent) before proceeding to the RNA-Seq protocol. RNA was processed as follows: 100 ng of total RNA were used to generate barcoded RNA-Seq libraries using the NEBNext Ultra II Directional RNA Library Prep Kit (New England Biolabs Inc). Libraries were sequenced on a NextSeq 2000 (Illumina) to generate 100 bases single-end reads. FastQ files for each sample were obtained using bcl2fastq v2.20.0.422 software (Illumina). The Next-generation sequencing (NGS) experiments were performed in the Genomics Unit of Centro Nacional de Investigaciones Cardiovasculares Carlos III (CNIC).

### Data processing, differential gene expression and gene set enrichment analysis

Single-end reads were quality controlled by removing adapter sequences, low-quality bases and short reads with FastQC (https://www.bioinformatics. babraham.ac.uk/projects/fastqc/) and Trim Galore (https://www.bioinformatics.babraham.ac.uk/projects/trim_galore/). Reads were aligned to the Ensembl human (hg38) reference genome annotation, version 109, using STAR [78], and quantification of expression levels were estimated using RSEM [79]. SAMtools [80] was used to sort, index, generate statistics and merge Binary Alignment Map (BAM) files. Post-alignment quality metrics were evaluated using RSeQC [81], Qualimap [82], dupRadar R package [83], Picard (http://broadinstitute.github.io/picard/), and MultiQC [84]. Gene level read count data was compared using R (v4.2.3) and the DESeq2 R package (v1.38.3) [85]. We utilized DESeq2 to normalize and correct the data by sequencing depth of the libraries and to test for differential expression, only genes that have a minimum of 10 counts are considered. Significant differentially expressed genes (DEG) were determined using a Benjamini-Hochberg false discovery rate (FDR) procedure [86], adjusted p-value <0.05. To remove noise, while preserving large differences and to control for small sample sizes, shrinkage of log2 fold change (LFC) was done using lfcShrink with apeglm method [87]. DEGs were considered if adjusted p-value < 0.05 and |LFC| >=1. We extracted the s-values [88] to rank the genes for a Gene Set Enrichment Analysis (GSEA). Enrichment analysis was performed using the ranked s-values through msigdbr (v7.5.1) (https://CRAN.R-project.org/package=msigdbr), clusterprofiler (v4.6.2) [89], and HTSanalyzeR2 (v0.99.19) R packages (https://github.com/CityUHK-CompBio/HTSanalyzeR2) using 10000 permutations to calculate statistical significance, a minimun geneset size of 10, and requiring a Benjamini-Hochberg FDR adjusted p-value <0.05 for statistical significance. Enrichment maps to visualize gene sets and Jaccard similarity coefficients between gene sets [90]. Gene sets used in this study were obtained from Reactome [91], (27 gene sets) and Kyoto Encyclopedia of Genes and Genomes [92] (43 gene sets) curated gene sets. Heatmaps were generated based on the Z-score values using the pheatmap R package (v1.0.12) (https://CRAN.R-project.org/package=pheatmap). Volcano, and point plots were generated with the ggplot2 R package (v3.4.1) (https://ggplot2-book.org/). Graphically, the significance of each pathway, according to the gene’s LFC, is represented using a circular visualization with circlize (v0.4.15) R package [93]. Finally, dplyr (v1.1.1) (https://CRAN.R-project.org/package=dplyr) and tibble (v3.2.1)(https://CRAN.R-project.org/package=tibble) R packages for efficient data manipulation.

### Detection of conditioned media-derived cytokines using antibody arrays

The Proteome Profiler Human XL Cytokine Array Kit (Bio-Techne R&D Systems) was used according to the manufacturer’s instructions. The conditioned media (CM) was concentrated using the Amicon® Ultra Centrifugal filters with nominal molecular weight limits (NMWLs) of 3 kDa. Briefly, each array was blocked and then, incubated overnight at 4 °C with 1 mL of concentrated CM from the different cell lines. After five washes, the array membrane was incubated with biotin-conjugated antibodies (1:250) for 1 h at RT, followed by an incubation with an HRP-linked secondary antibody (1:1000) for 1 h at RT. Membranes were incubated with a chemiluminescent substrate and imaged within 15-25 min to avoid chemiluminescent signal fading.

### Comet assay

Cells were cultured in complete media and digested using trypsin for 3 min. Then, cells were collected and spun at 300 x g for 5 min. The medium was aspirated and the cells were suspended at 1 x 10^6^ cell/mL in complete media. The cell suspension was combined with 1% molten low melting point agarose (at 37 °C) at a ratio of 1:10 (v/v), mixed gently by pipetting up and down, and immediately pipetted onto a slide. The slide was placed flat at 4 °C for 10 min. The slides were then immersed in 2.5 M NaCl, 100 mM disodium EDTA, 10 mM Tris base, and 200 mM NaOH (Lysis buffer). The slides were gently removed from the Lysis buffer and were immersed in 200 mM NaOH and 1 mM disodium EDTA (alkaline electrophoresis solution), pH >13 for 1 h to allow DNA unwinding. Pre-chilled alkaline electrophoresis solution was added in the electrophoresis slide tray, not exceeding 0.5 cm above the slides. The slides were placed inside and covered with a cap. The power supply voltage was set to 1 V/cm and run for 30 min. Excess electrophoresis solution was drained from the slide and 50 - 100 µL of propidium iodide staining solution was placed onto each dried agarose and stained for 5 min at RT in the dark. The slides were briefly rinsed in dH2O and dried completely at 37 °C in the dark. The images were acquired and analyzed. Tail moment was measured with the OpenComet plug-in from ImageJ software. The comets that were wrongly recognized by the program were removed manually.

### Statistical analysis

Statistical analysis was performed with GraphPad Prism software (version 7.00 and 8.00). Data is represented as mean ± S.E.M. Differences between two groups were estimated with two-tailed unpaired Student’s *t* test or the Mann-Whiney U-test. A minimum sample size of 3 independent experiments was used in all experiments for statistical comparisons. *P* value < 0.05 was considered statistically significant. \**P*< 0.05, \*\**P< 0.01, ***P*< 0.0001.

---

**Supplementary Figure 1.**
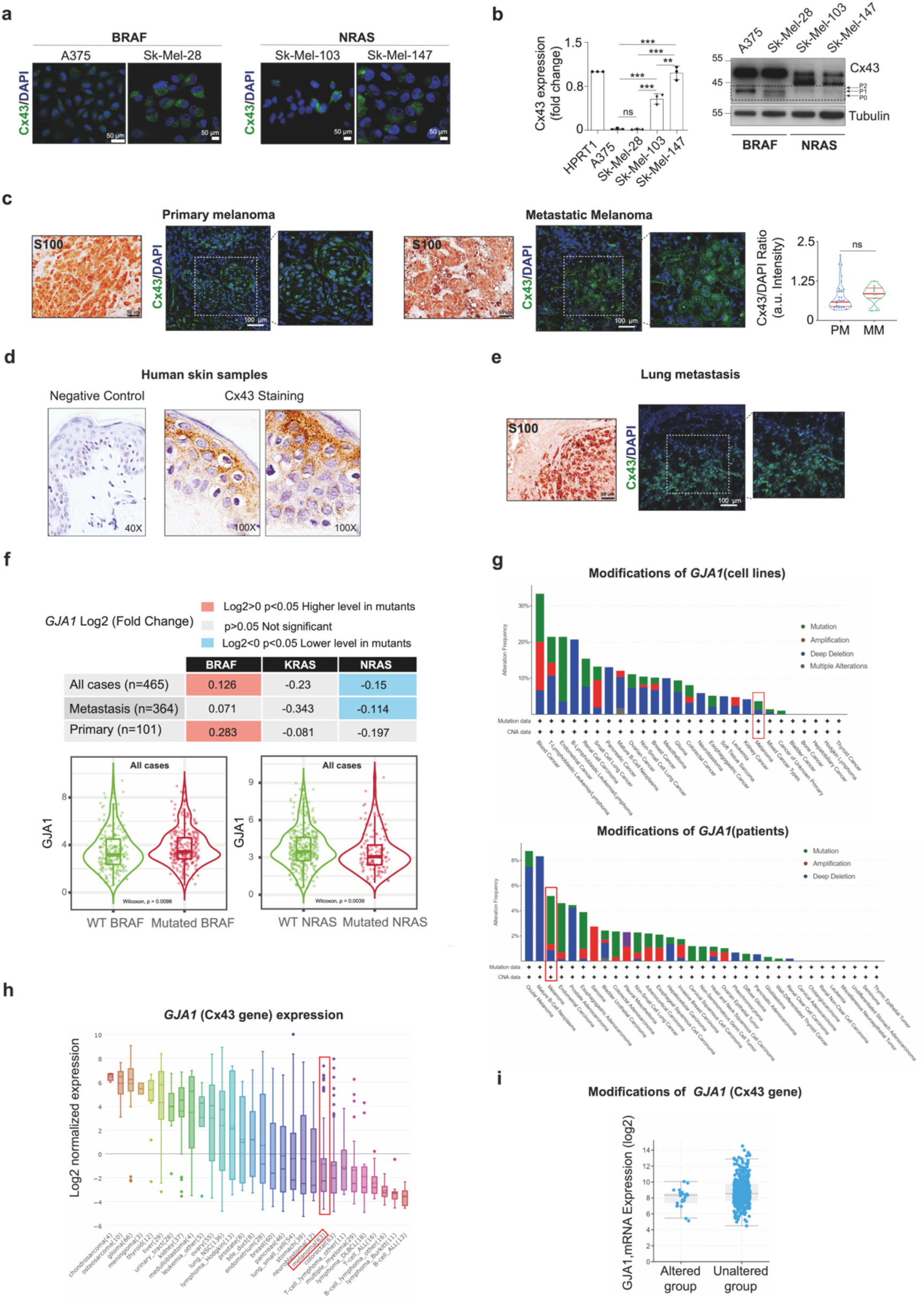
(a) Confocal images of Cx43 expression (green) in different melanoma cell lines with BRAF (A375 and Sk-Mel-28) and NRAS (Sk-Mel-103 and Sk-Mel-147) mutation. Scale bars: 50 µm (Cx43 and DAPI). (b) Cx43 mRNA (left) and protein (right) levels in BRAF and NRAS melanoma cell lines. (c) Confocal images and quantification of Cx43 expression (green) in primary (n=19) and metastatic melanoma (n=4). DAPI (blue) was used for nuclear staining. Staining of the melanocytic marker S100. Scale bars: 50 µm (S100) and 100 µm (Cx43 and DAPI). (d) Images of Cx43 IHQ staining in human skin epidermis. (e) Confocal images of Cx43 expression (green) in lung metastasis. DAPI (blue) was used for nuclear staining. S100 staining of the melanocytic marker S100. Scale bars: 50 µm (S100) and 100 µm (Cx43 and DAPI). (f) (e) GJA1 alteration frequency (mutation, amplification, deep deletion and multiple alterations) in Cancer Cell Line Encyclopedia with 1,739 tumour cell line samples (CCCLE, Broad 2019) and in NCI-60 Cell Lines with 67 tumour cell line samples (NCI, Cancer Res 2012). GJA1 alteration frequency based on TCCA PanCancer Atlas Studies (10,953 patients/10,967 tumour samples). Images from www.cbioportal.org. (g) GJA1 expression from Illumina RNASeq (log2) in altered and unaltered groups, p-Value 6.64e-3. (h,i) On the top, association of GJA1 gene mutation levels represented in LFC, comparing BRAF, KRAS and NRAS mutation status of melanoma samples, according to the TIMER database (v. 2; http://timer.cistrome.org/). On the bottom, association of GJA1 gene expression with BRAF and NRAS mutation status of melanoma samples (clustered cases and metastasis), according to the TIMER database (v. 2; http://timer.cistrome.org/).

**Supplementary Figure 2.**
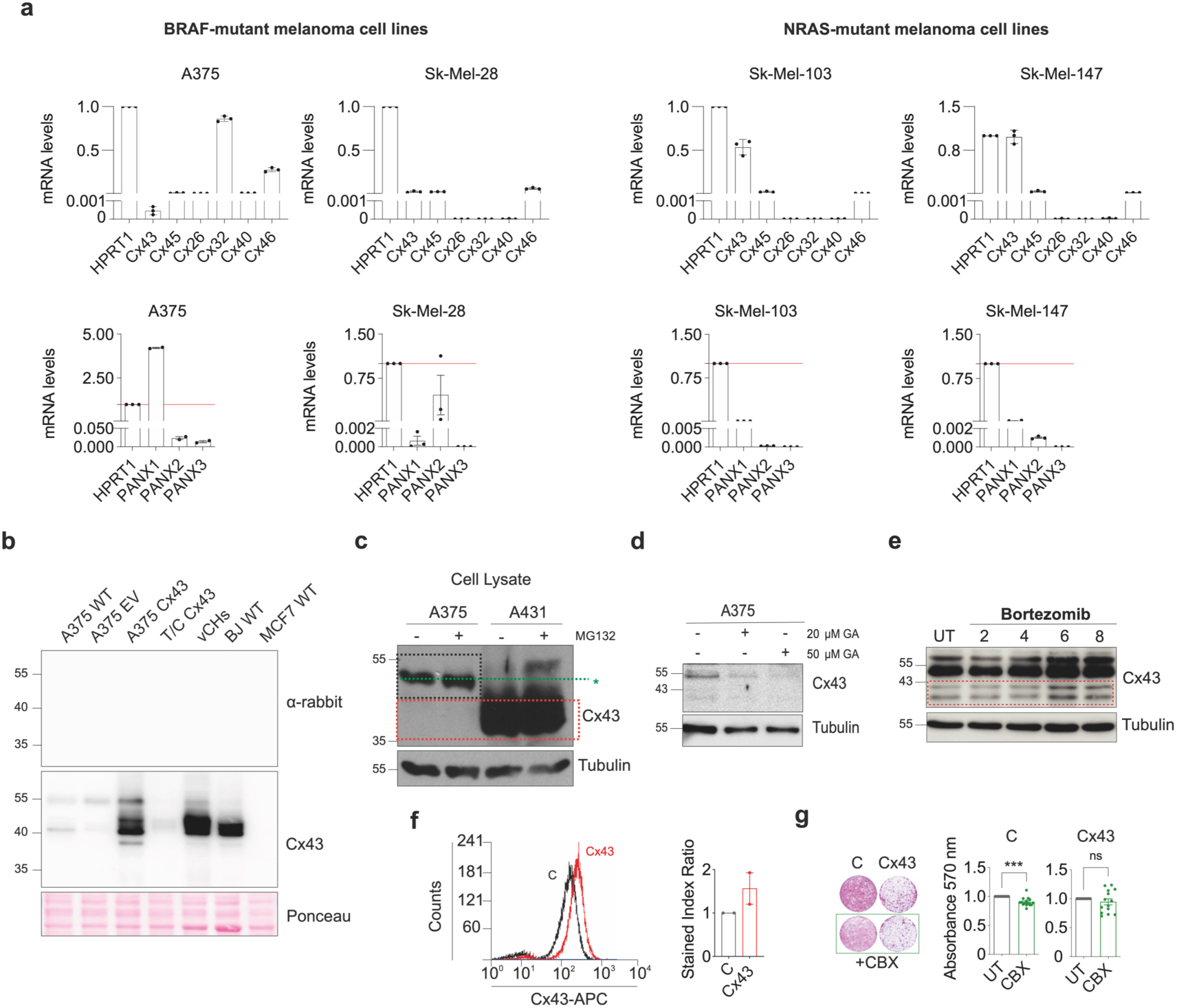
(a) RT-qPCR of different connexins and pannexins in BRAF and NRAS human melanoma cell lines (A375, Sk-Mel-28, Sk-Mel-103 and Sk-Mel-147) (n=3). (b) Immunoblotting of Cx43 in different human cell lines. Human melanoma cell lines (A375-C and A375 Cx43), chondrocyte cell line (T/C 28a2),a primary vertebral chondrocytes (vCH), fibroblast cell line (BJ) and a breast cancer cell line (MCF7). Negative control of secondary antibody (anti-rabbit) is shown. Ponceau was used as loading control. (c) Immunoblotting of Cx43 under the treatment of the proteasome inhibitor, MG132 in human melanoma (A375) and human carcinoma epidermoid (A431) cell lines. Tubulin was used as loading control. (d) Immunoblotting of Cx43 under two concentrations of the SUMOylation inhibitor, ginkgolic acid (GA), in A375 cell line. Tubulin was used as loading control. (e) Immunoblotting of Cx43 under the treatment of proteasome inhibitor, bortezomib (BTZ), in A375 cell line at different times. (f) Flow cytometry of Cx43 stained index in A375-C and A375-Cx43 cell lines. (g) Images and quantification of the colony formation capacity of melanoma cells treated with 100 µM of the hemichannel blocker, carbenoxolone (CBX), for 7 days in A37-C and A375-Cx43 cell lines (n= 3, with 5 replicates). Two-tailed Student’s t-test were used to calculate the significance represented as followed: ns: no significant, * p < 0.05, ** p < 0.01. *** p < 0.001. Data is presented as mean ± SEM.

**Supplementary Figure 3.**
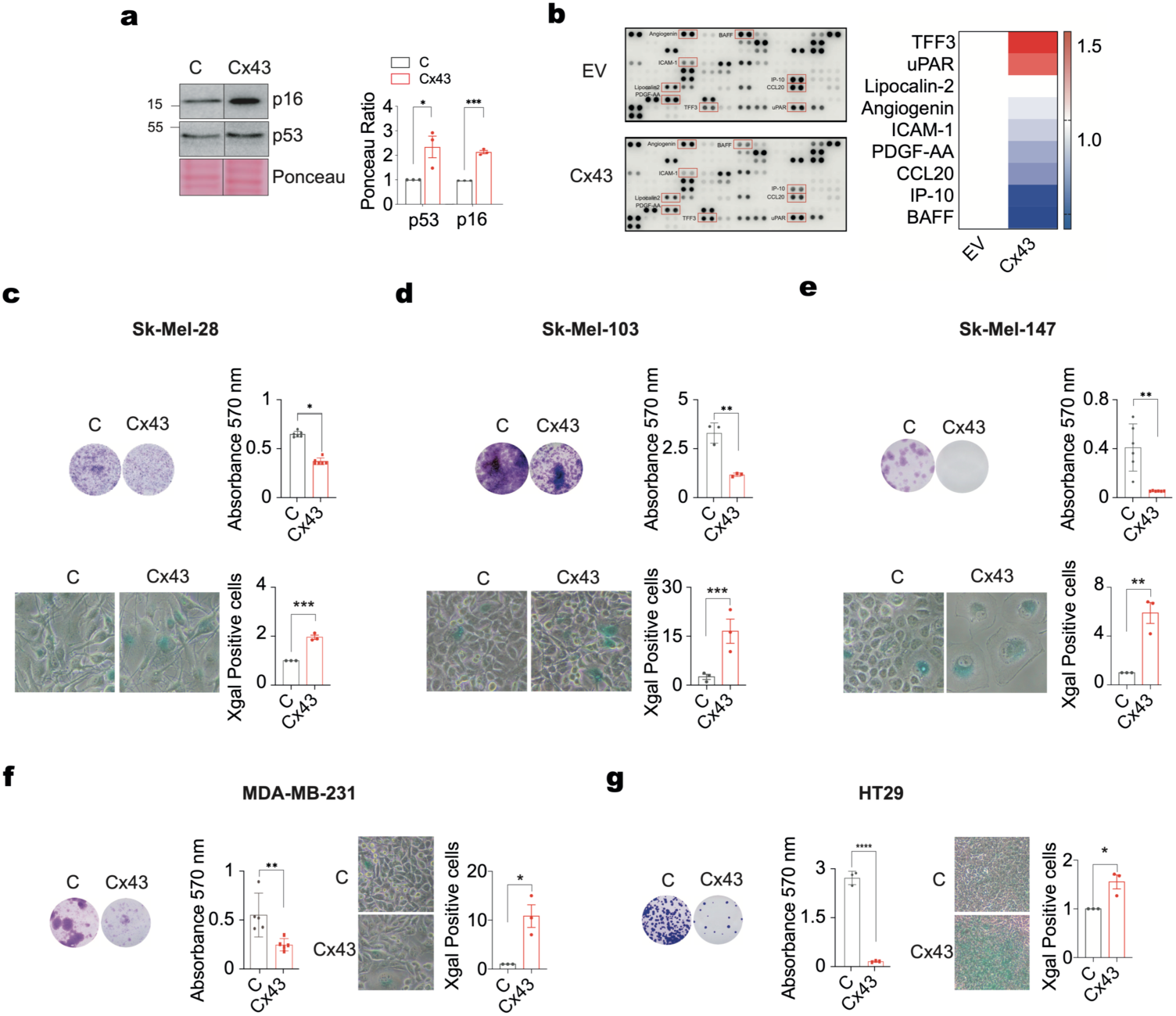
(a) Immunoblotting and quantification of the senescence marker p53 and p16 (n= 3). Ponceau was used as loading control. (b) Heatmap and immunoblotting of cytokines arrays in A375-C and A375 Cx43 cell lines. (c-g) Staining, images and quantification of colony formation assays in BRAF (Sk-Mel28, MDA-MB-231 and HT19) and NRAS (Sk-Mel103, Sk-Mel147 and BLM) mutant cell lines under the overexpression of Cx43. Representative images of SAß-Galactosidase activity and quantification (n= 3) in BRAF and NRAS mutant cell lines under the overexpression of Cx43. Two-tailed Student’s t-test were used to calculate the significance represented as followed: ns: no significant, * p < 0.05, ** p < 0.01. *** p < 0.001. Data is presented as mean ± SEM.

**Supplementary Figure 4.**
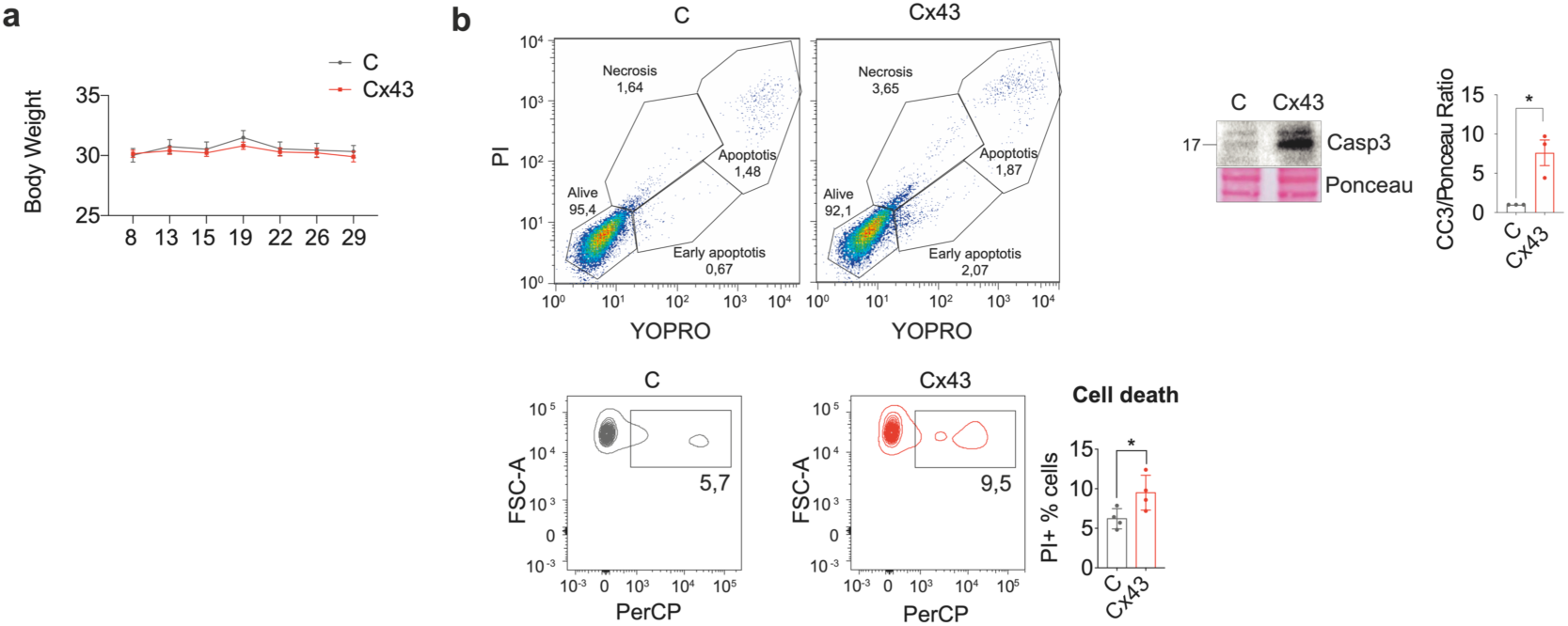
(a) Body weight remains unaffected in nude mice injected with A375-C and A375-Cx43 cells at different time points throughout the experiment. (b) Cell death in A375-C and A375-Cx43 measured by flow cytometry (n=4) and western-blot of CC3 levels.

**Supplementary figure 5.**
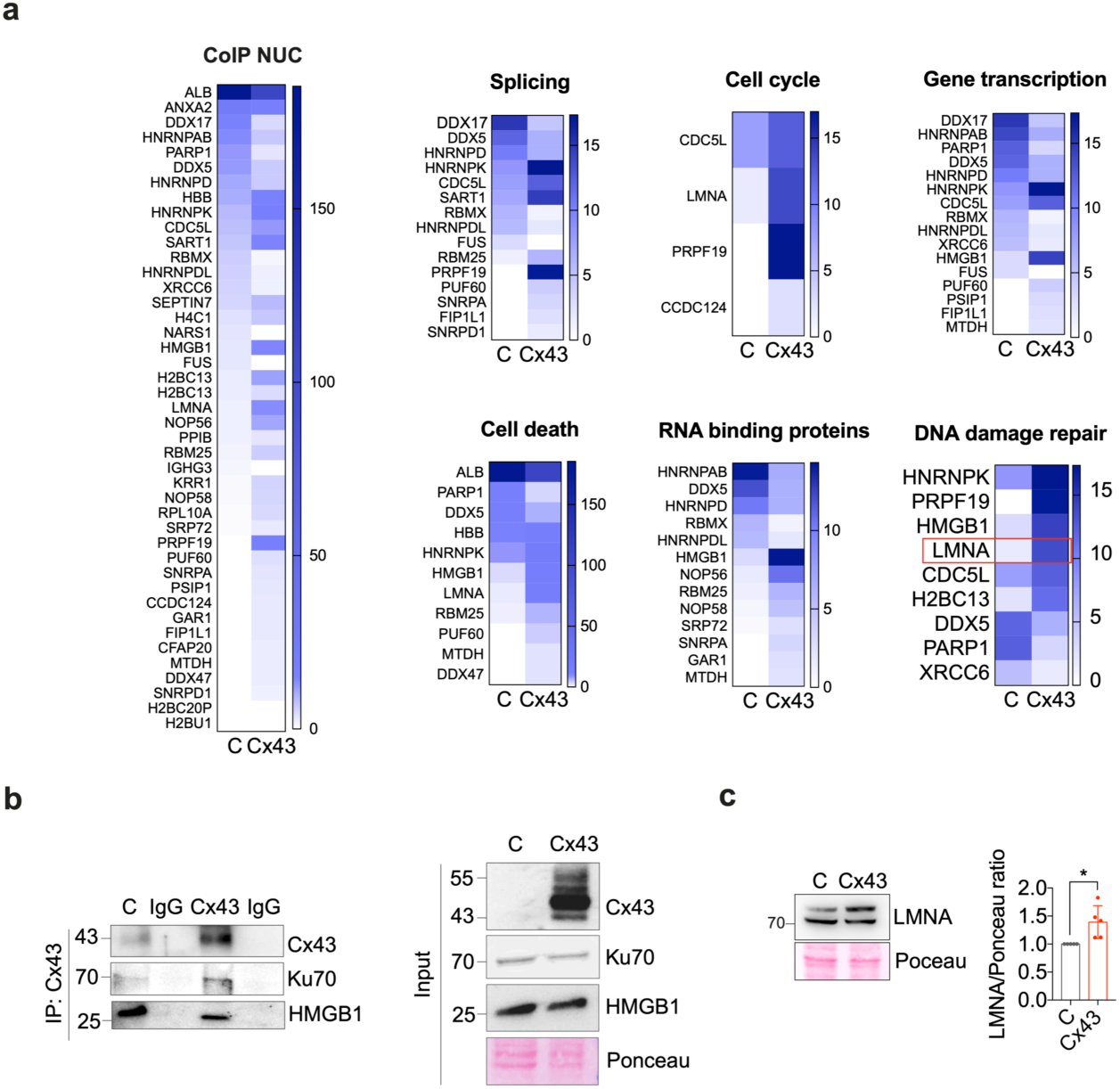
(a) Heatmaps representing the spectral count values and the −10log p-value of Cx43 interactor proteins in the nucleus of A375-C and Cx43 cells. Those proteins are involved in splicing, cell cycle, gene transcription, cell death and RNA binding processes among others, showing a differential expression pattern between both groups. (b) Cx43 IP NUC of A375-C and Cx43 cells. Input is shown on the right, while IP is shown on the left. IgG is indicated as a negative control. (c) LMNA protein levels measured by western-blot.

**Supplementary figure 6.**
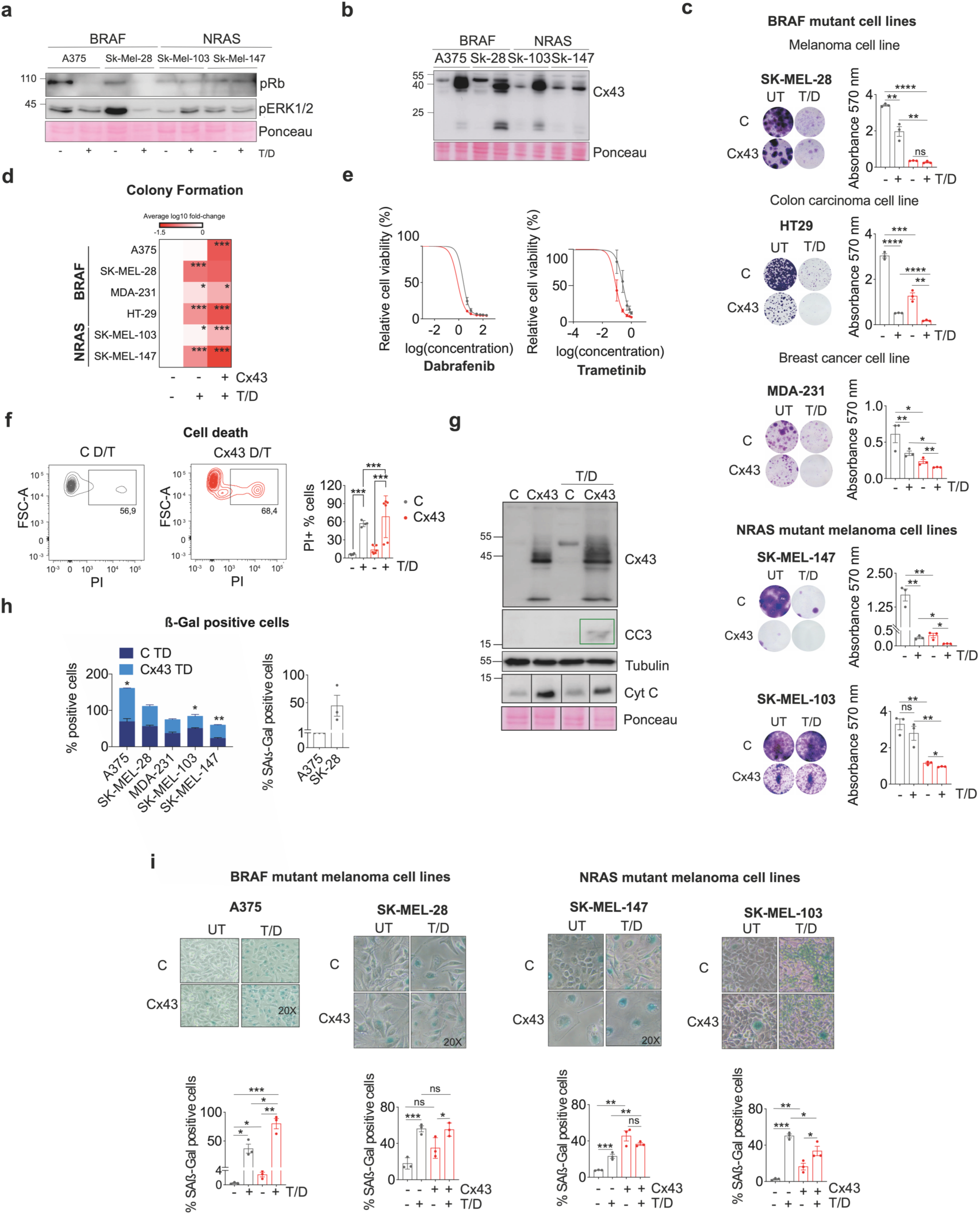
(a) Protein levels of pRB and pERK1/2 in BRAF (A375, Sk-Mel-28) and NRAS (Sk-Mel-103, Sk-Mel-147) mutated cell lines, in the presence/absence of trametinib (1 nM) and dabrafenib (100 nM) for 72 h. **(b)** Cx43 protein levels and pattern of BRAF (A375, Sk-Mel-28) and NRAS (Sk-Mel-103, Sk-Mel-147) mutated cell lines, in the presence/absence of trametinib (1 nM) and dabrafenib (100 nM) for 72 h **(c)** Colony formation assay of BRAF (Sk-Mel-28, HT29, MDA-231) and NRAS (Sk-Mel-103, Sk-Mel-147) mutated cell lines, in the presence/absence of Trametinib (0,1 nM) and Dabrafenib (10 nM) for 1 week. **(d)** Heatmap showing the quantifications of the colony formation assay showed in (**c**). Data is represented as the average of LFC. **(e)** BRAF/MEKi showed significant cytotoxicity against the A375-Cx43 cell line (dabrafenib IC50 = 0.7782 nM and trametinib IC50 = 0.0803 nM) compared to A375-C (dabrafenib IC50 = 2.301 and trametinib IC50 = 0.2577 nM). **(f)** Flow cytometry of cell death in A375 EV and A375 Cx43 cells in the presence of BRAF/MEKi (1 nM Trametinib/100 nM dabrafenib) for 72 h. Data represents PI positive cells. **(g)** Western-blot showing an increment in CC3 and CytC levels in the presence of Cx43 and under trametinib and dabrafenib (T/D) for 72h. **(i)** Representative images of SA-ß-Gal positive cells of BRAF (A375, Sk-Mel-28, MDA-231) and NRAS (Sk-Mel-103, Sk-Mel-147) mutated cells, overexpressing Cx43 and treated with BRAF/MEKi (TD) for 72 h. **(h)** Comparative histogram of SAß-Galactosidase activity in BRAF and NRAS melanoma cell lines treated with combination of 100D/1T for 5 days, in presence and absence of Cx43. All experiments were performed in at least three independent replicates. **(i)** Treatment of BRAF (A375, Sk-Mel-28) and NRAS (Sk-Mel-147, Sk-Mel-103) mutated cell lines with BRAF/MEKi (TD) in the presence/absence of Cx43 increases the number of senescence-associated β-galactosidase positive cells. All experiments were performed at least three independent times. Data is presented as mean ± SEM. Two-tailed Student’s t-test and one-way ANOVA were used to calculate the significance represented as followed: * P < 0.05, **P < 0.01, ***P < 0.0001.

**Supplementary figure 7.**
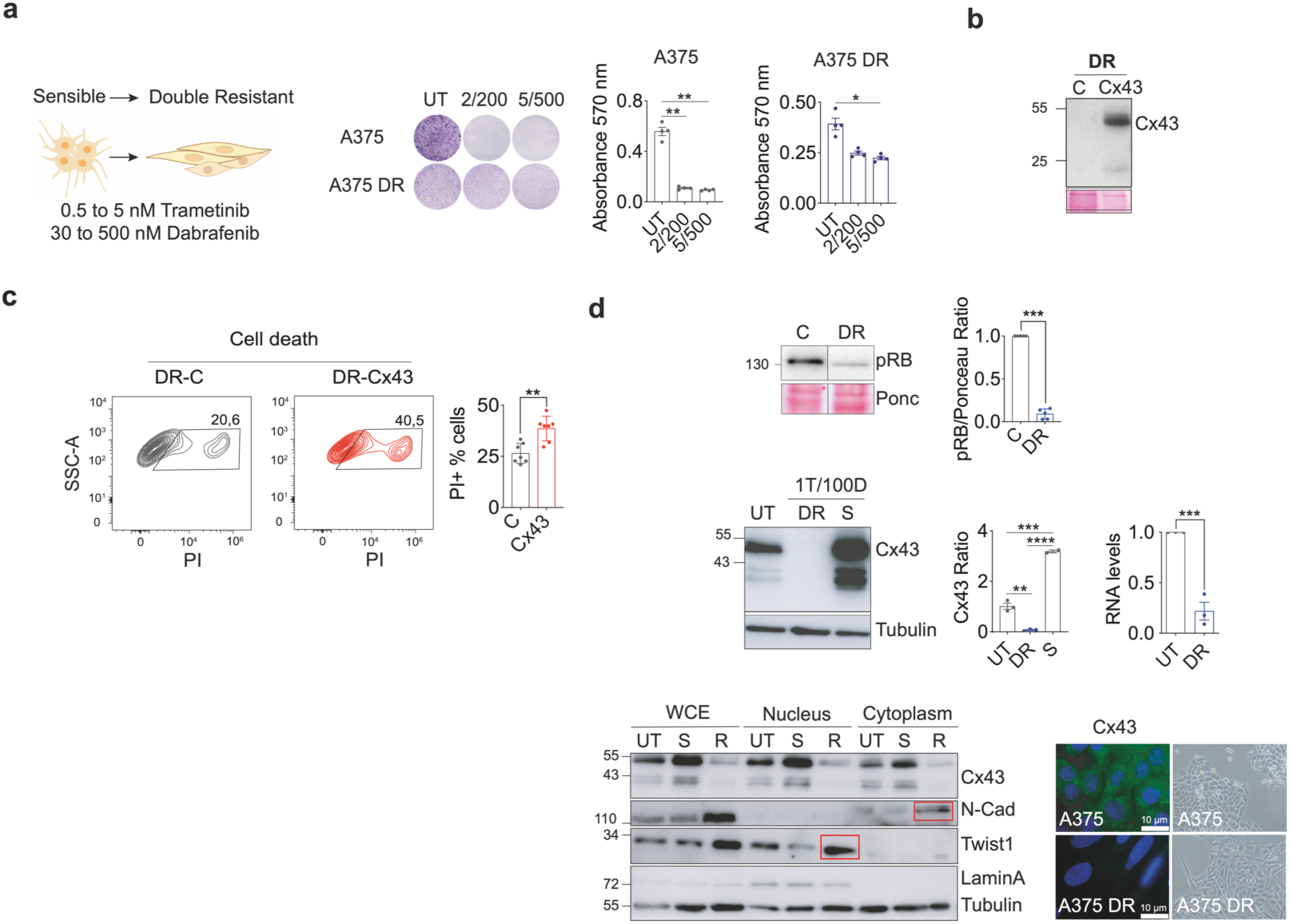
(a) A375 melanoma cells become resistant to high concentrations of BRAF/MEKi (5 nM Trametinib/500 nM Dabrafenib). Colony formation assay showing A375 DR cells are not affected by BRAF/MEKi compared to parental A375 cells (WT). (B) immunoblotting of DR cells transfected with Cx43 (DR-Cx43) and its empty vector counterpart. **(c)** Flow cytometry of cell death in A375 DR overexpressing Cx43 in the presence of BRAF/MEKi. Data represents PI positive cells detected by PerCP laser (n=3) **(d)** On the top, A375 DR cells show lower levels of pRB protein, as well as a lack of Cx43 protein and mRNA levels compared to cells that respond to the treatment (S). On the bottom, epithelial to mesenchymal transition markers are regulated in resistant cells, while N-Cadherin (N-Cad) is upregulated in the cytoplasm fraction, Twist1 appears upregulated in the nuclear fraction, compared to untreated (UT) and sensible (S) cells. LaminA and Tubulin are used as nucleus/cytoplasm extraction quality control, as well as loading control. Data is presented as mean ± SEM. Two-tailed Student’s t-test and one-way ANOVA were used to calculate the significance represented as followed: * P < 0.05, **P < 0.01, ***P < 0.0001.

